# A toolkit for mapping cell identities in relation to neighbours reveals Notch-dependent heterogeneity within neuromesodermal progenitor populations

**DOI:** 10.1101/2024.09.03.610492

**Authors:** Matthew French, Rosa Portero, J. Kim Dale, Guillaume Blin, Val Wilson, Sally Lowell

**Author notes:** co-senior authors (Guillaume Blin, Val Wilson, Sally Lowell).

## Abstract

Patterning of cell fates is central to embryonic development, tissue homeostasis, and disease. Quantitative analysis of patterning reveals the logic by which cell-cell interactions orchestrate changes in cell fate. However, it is challenging to quantify patterning when graded changes in identity occur over complex 4D trajectories, or where different cell states are intermingled. Furthermore, comparing patterns across multiple individual embryos, tissues, or organoids is difficult because these often vary in shape and size.

Here we present a toolkit of computational approaches to tackle these problems. These strategies are based on measuring properties of each cell in relation to the properties of its neighbours to quantify patterning, and on using embryonic landmarks in order to compare these patterns between embryos. We use this toolkit to characterise patterning of cell identities within the caudal lateral epiblast of E8.5 embryos, revealing local patterning in emergence of early mesoderm cells that is sensitive to inhibition of Notch activity.

## Introduction

Quantifying patterning at single cell resolution establishes when and where cell fate decisions are occurring, and helps clarify mechanisms by which cells make differentiation decisions. Impressive progress has been made in measuring and interpreting patterning of cell identity in tissues that exhibit relatively simple graded or striped patterns that can be readily visualised and quantified in 2D [1]. However, in many biological systems, cells change identity in relatively complex patterns across curved 3D shapes, making it challenging to quantify these patterning events.

In recent years, improvements to segmentation have made it possible to measure marker expression at single cell resolution within intact 3D tissues [2–7]. This opens up new opportunities for using neighbour-relationships to quantify patterning in 3D [8,9]. For example, measuring differences between neighbours should make it possible to map complex graded changes across 3D space. Additionally, measuring local heterogeneity in cell identity provides clues about the logic of local interactions that coordinate cell fate changes. For example, particular types of spotty pattern can be generated by lateral inhibition [10] or stochastic cell fate allocation [11], while locally-coherent cell identities are consistent with lateral induction [12] homeogenetic induction [13], quorum-sensing [14] or community effects [15–17]. Measuring changes in patterning after experimental manipulation of candidate regulators could then be used to identify mechanisms underlying local coordination of cell fate.

Approaches to measure patterning in a number of contexts have been developed [18–28]. However, challenges remain in measuring mesoscale patterning in cell identity during development [29]. This is in part because individual embryos differ from one another in size or shape, so there is a need for unbiased methods to define and normalise regions of interest that can be compared between different embryos [30,31]. Furthermore, defining different cell states, and relating the state of one cell to the states of its neighbours, can become particularly challenging when different cell identities merge along a continuum rather than falling naturally into discrete states. These problems are exemplified by the axial progenitor region in mouse embryos, which forms a complex curved surface with graded expression of known regulators of cell lineage, and has a well-defined fate map.

The axial progenitor region in E8.5 mouse embryos encompasses the caudal epiblast, which contains neuromesodermal progenitors (NMPs). These are bipotent progenitors that can generate neuroectoderm (producing spinal cord) and paraxial mesoderm (producing musculoskeleton) during axis elongation [32]. Cells exit from the NMP region in two ways. From the anterior edge of the NMP region they commit to neuroectoderm differentiation. Alternatively, they move towards the midline primitive streak, where they are committed to mesoderm differentiation, undergoing epithelial-to-mesenchymal transition and exiting towards the paraxial mesoderm. Differentiation of NMPs is governed by the activity of Wnt and FGF [32,33], but local cell-cell heterogeneity in cell identity [34,35] suggests that locally-acting feedback signalling between close neighbours may also contribute to controlling changes in NMP identity.

Here, we focus on the transition of NMPs towards a mesodermal fate, taking advantage of TBX6 as an early marker of mesoderm commitment [36–38]. By analysing neighbour relationships, we identify steep local changes in TBX6 expression in the medial regions of the NMP progenitor region and show that TBX6 positive cells in this region tend to be surrounded by TBX6-negative cells and organised in a non-random distribution.

Notch signalling coordinates cell fate decisions between neighbouring cells in many contexts, often acting by lateral inhibition [10] to generate local heterogeneity in cell identity but sometimes mediating lateral induction [12] to ensure local coherence of cell identity. Notch signalling favours mesoderm differentiation from hESC-derived NMPs at the expense of neural fate [39], positioning Notch as a likely candidate for generating heterogeneity in cell identity as NMPs differentiate into mesoderm during axis elongation. To test this, we extended our patterning analysis to embryos that had been subjected to Notch inhibition and found that local heterogeneity of TBX6 depends on Notch signalling.

We present a generalisable toolkit for describing local patterning of cell identity and for revealing phenotypic changes in response to candidate regulators. We demonstrate their utility in exploring patterning of neuromesodermal progenitors (NMPs) and their differentiated derivatives during axis elongation in mouse embryos and gastruloids.

## Results

### Challenges for unbiased quantification of patterning

Neuromesodermal progenitors in the node-streak border and caudolateral epiblast are marked by coexpression of TBXT and SOX2 [32]. As NMPs commit to a mesoderm fate, they upregulate TBX6 [36–38]. Therefore, the question of when and where NMPs become mesoderm-committed should in principle be answered by staining embryos for these three markers and analysing the resulting images.

However, there are a number of challenges in interpreting such images. For example, TBXT and SOX2 exhibit graded distributions (Figure 1A,B), and their gradients do not appear to be aligned to a single axis (Figure 1A,B). This makes it challenging to objectively define an ‘NMP containing region’ based on visual inspection alone. Furthermore, the mesoderm commitment marker TBX6 does not seem to be upregulated at a clear boundary, but rather is expressed in a subset of cells within the caudolateral epiblast (Figure 1A,B: note contrast with the seemingly homogenous expression in the presomitic mesoderm). It is not easy to determine by eye whether these sporadic TBX6+ cells are restricted to a particular location or expressed in a particular pattern.

**Figure 1:**
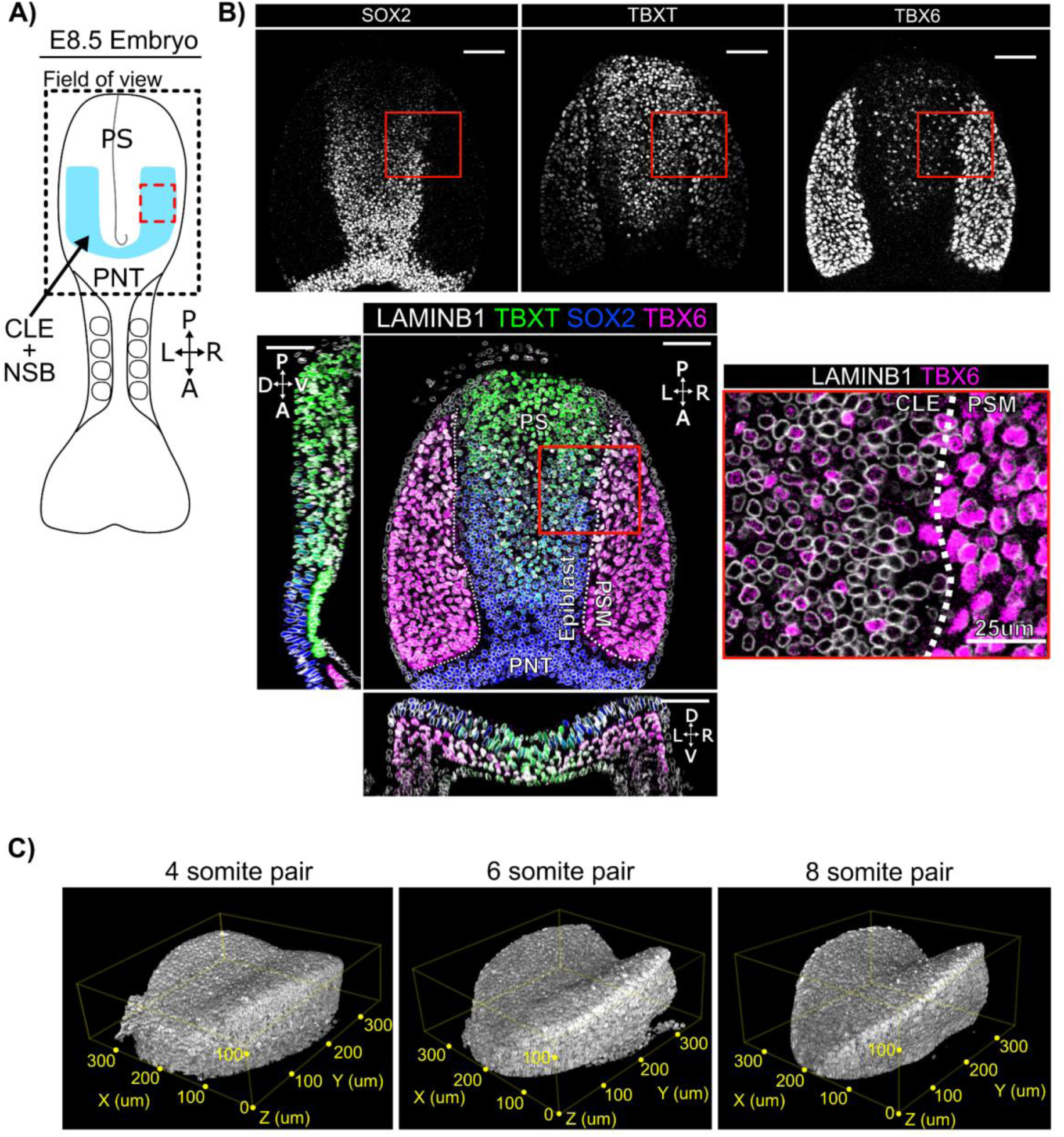
Spatial patterning of TBXT, SOX2 and TBX6 in the posterior E8.5 mouse embryo A: Diagram representing a 4-somite pair E8.5 embryo and showing the positioning of the confocal imaging field of view (FOV) used in b and c as a dashed outline. The FOV encompasses the primitive streak (PS), and a domain comprising the neural & mesodermal bi-fated caudal lateral epiblast (CLE) plus node streak border (NSB) in blue. Anterior (A)/ Posterior (P) and Left (L)/Right (R) and posterior neural tube (PNT) are indicated. The dotted red box corresponds approximately to the insert shown in (b). B: Confocal z-slices across each plane of the field of view shown in (a) with insert (red box) showing the spatially heterogeneous TBX6 expression in the SOX2+TBXT+ CLE, and the spatially homogeneous TBX6 expression in PSM. White dotted line indicates the boundary between epiblast and the presomitic mesoderm (PSM). Unless specified, scale bars indicate 50 µm. C: 3D renders of LaminB1 signal at 4, 6, and 8 somite pair stage E8 embryos. Highlighting the increase in ‘Pringle-like’ epiblast curvature and the associated challenge posed when assessing patterning in 3D.

Additional challenges arise from the fact that embryos comprise multiple tissues arranged in 3D space: this can create confusion when embryos are viewed in 2D. In particular, the caudal epiblast (the focus of this study) lies on top of newly-formed mesoderm and has a complex curved 3D shape, making it difficult to visualise and quantify patterning in and around this region in a 2D representation (Fig1A,B). Finally, there is the problem that different embryos have slightly different shapes and sizes, so it is challenging to compare quantitative measures of patterning between embryos. This becomes particularly important when assessing reproducibility of particular patterns, or when assessing patterning phenotypes after experimental manipulation of candidate regulators.

To address these problems, we sought quantitative computational methods to:

- Facilitate faithful 2D visualisation of 3D patterning
- Visualise averaged patterning across multiple embryos of the same developmental stage
- Define regions of interest in a way that avoids arbitrary thresholding of individual markers
- Use neighbour-relationships to quantify and visualise the shape and steepness of gradients.
- Use neighbour-relationships to quantify and visualise fine-grained patterning.
- Compare patterning between different experimental conditions.

### 3D Epiblast projection and alignment method: PRINGLE

Our first aim was to facilitate faithful visualisation of 3D patterning in 2D, and to enable comparison of patterning between embryos. To achieve this, we developed a pipeline to computationally flatten the complex curved epiblast shape and to normalise the edges of the epiblast to defined landmarks.

We used a custom-written software package, PickCells, that combines tools for nuclear segmentation (Blin et al. 2019) with methods for quantifying the properties and positions of each cell within densely packed 3D tissues. In this pipeline, immunofluorescence confocal stacks are pre-processed to extract nuclear fluorescence intensity and neighbourhood information (Figure 2A). In brief, nuclei are segmented using LAMINB1 nuclear envelope staining [3] to identify individual nuclei and to quantify the fluorescence intensity for SOX2, TBXT, and TBX6 immunofluorescence. The epiblast is manually isolated using PickCells software, and the nearest neighbours of each nucleus are identified in 3D using Delaunay triangulation (see Methods).

**Figure 2.**
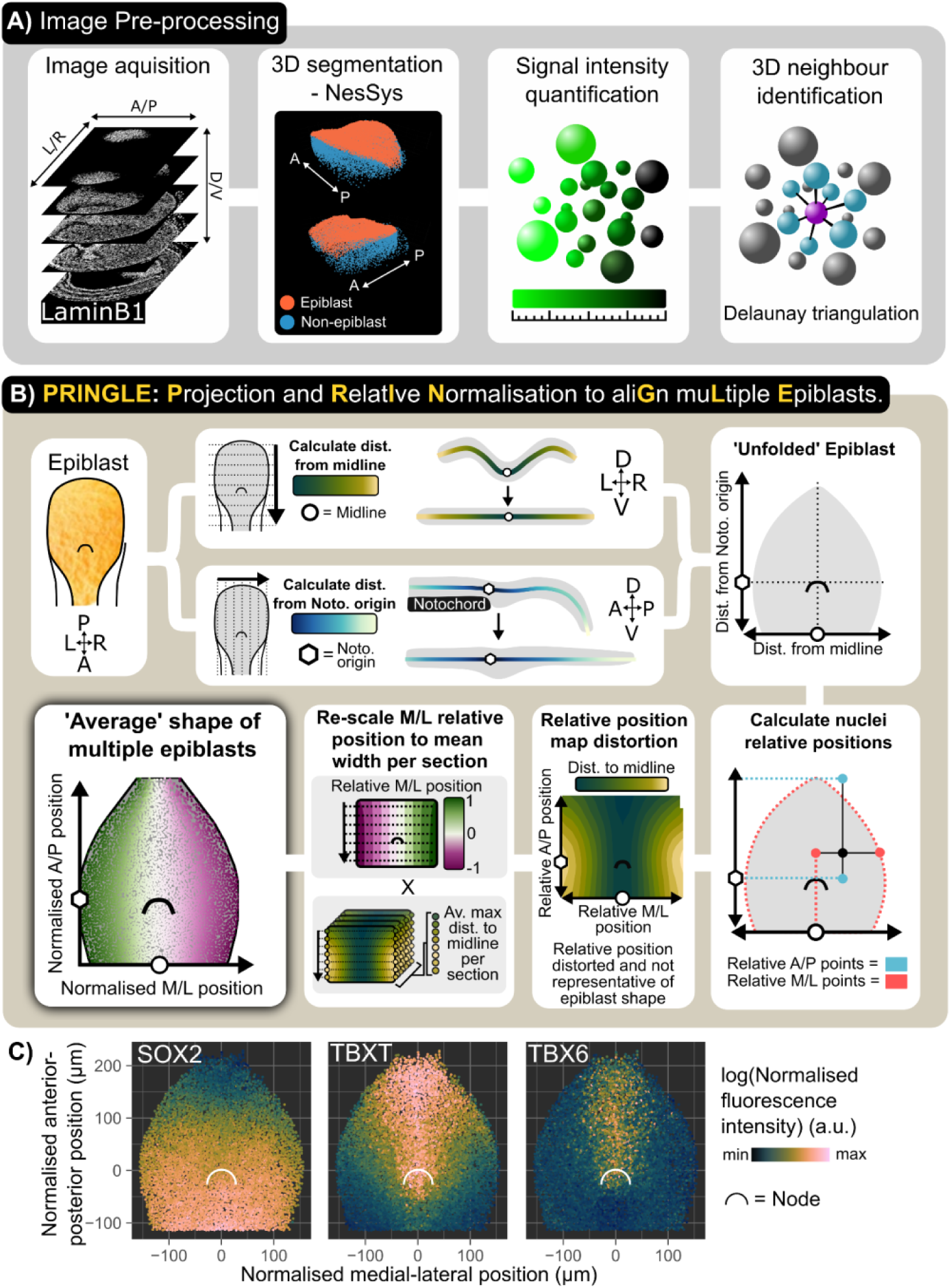
3D Epiblast projection and alignment method: PRINGLE. A: Single cell analysis pipeline: individual nuclei are identified from 3D confocal images and manually labelled as epiblast or non-epiblast using the PickCells software. Fluorescence intensity is quantified within each nucleus and the neighbours of each cell in 3D are identified. B: Summary of PRINGLE method to project the 3D nuclei centroids in 2D and align multiple embryos: The distance of nuclei centroids are calculated from the midline and notochord along principal curves in transverse and sagittal sections respectively. These distances are then used as coordinates to project nuclei in 2D and create an unfolded epiblast. Unfolded maps generated from multiple embryos are then registered together by normalising nuclei coordinates relative to tissue landmarks: the midline and edge of epiblast are used as landmarks for the left/right (L/R) axis, and the notochord origin and primitive streak posterior tip as landmarks for anterior/posterior (A/P) axis. However relative positions produce a distortion, therefore relative positions are normalised to the average epiblast width. This registration procedure results in an ‘average’ epiblast map in 2D. D: dorsal, V: ventral, A: Anterior, P: Posterior, M: Midline, L: Lateral edge of epiblast. This approach is explained in more detail in Supp Figure 1. C: A four-somite-pair epiblast showing SOX2, TBXT, and TBX6 expression (measured as outlined in (a)) mapped onto the average epiblast shape of multiple four-somite-pair epiblasts (established as outlined in (b)). Points represent nuclei centroids.

The epiblast is challenging to visualise in 2D because it curves in 3D convex and concave planes (similar to the shape of a Pringle potato crisp). We developed an algorithm which utilises Projection and Relative Normalisation to aliGn muLtiple Epiblast (PRINGLE) (Figure 2B, Supp Figure 1). This has the effect of computationally flattening the curved epiblast, and makes it possible to unify multiple embryos by using relative positions of nuclei to landmark tissue structures, including the midline, node (manually labelled (Figure 2B, Supp Figure 2), and epiblast edge (Figure 2B, Supp Figure 1). This provides absolute and relative positions of nuclei in relation to these landmarks. Using this approach, we visualised levels of SOX2, TBXT, and TBX6 immunofluorescence on individual PRINGLE projection epiblasts, as a starting point for further analysis (Figure 2C).

### Mapping the spatial distribution of Neuro-Mesodermal progenitors within the CLE

We next sought to map the NMP-containing region within and around the caudal-lateral epiblast as a starting point for asking when and where TBX6+ mesoderm arises in relation to this region. Grafting of microdissected regions [40,41] has identified areas of the CLE that contain cells with both neural and mesodermal potency (black boxes in Figure 3A). However, this functional approach does not fully define the NMP region: for example, NMPs may also exist in some closely surrounding regions that were not included in grafts.

**Figure 3.**
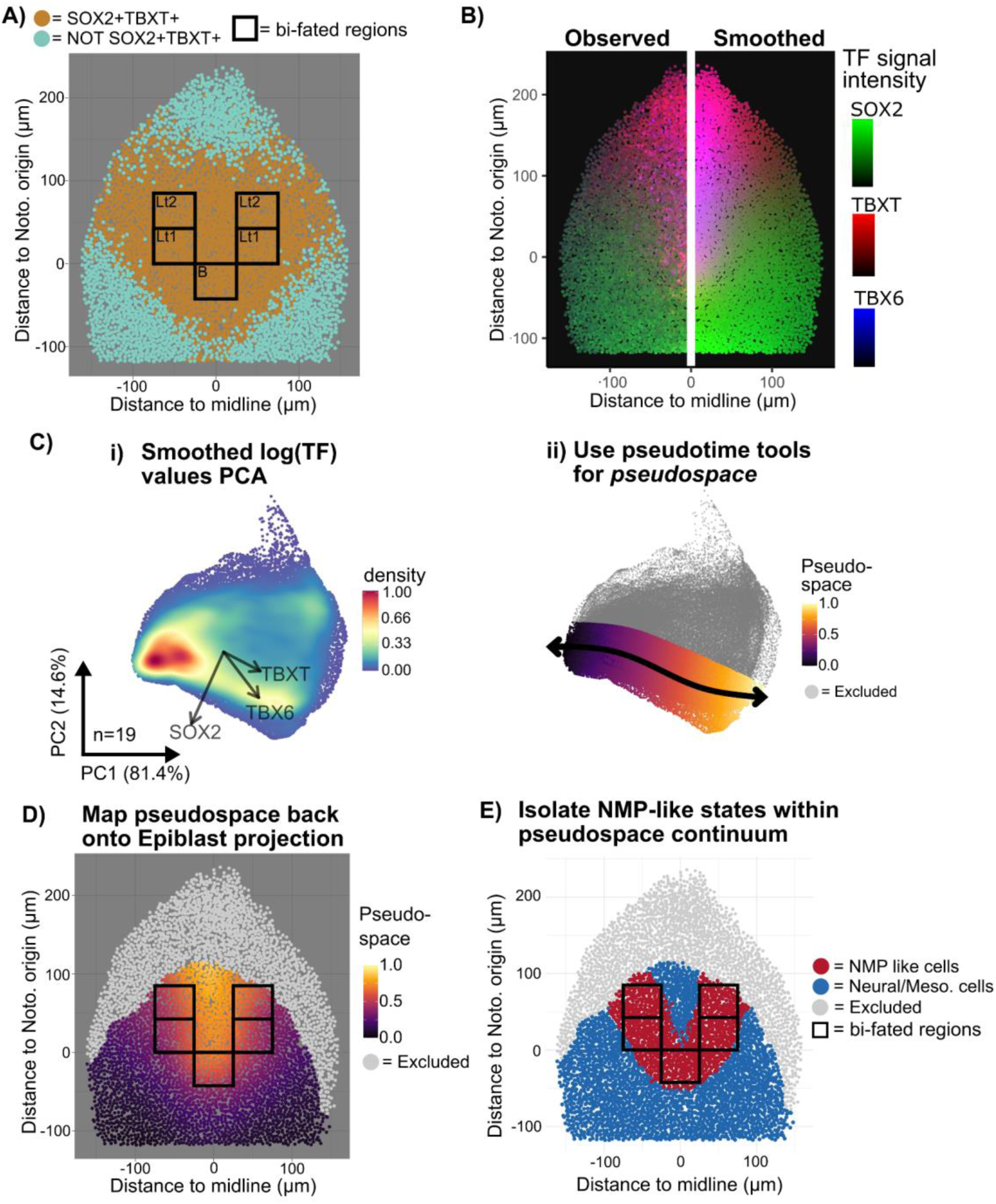
Mapping the spatial distribution of Neuro-Mesodermal Progenitors A: Manifold projection of nuclei centroids in an example 4-somite-pair embryo showing SOX2+TBXT+ cells (orange) and the regions in the lateral epiblast (Lt1-2) and node-streak border (B) defined by grafting experiments (add ref here) as having neural and mesodermal potency (bifated regions: black boxes). SOX2+ TBXT+ are not restricted to the known N-M-bi-fated regions and instead map onto a broader spatial region. B: Iterative kernel averaging spatially smooths TF intensity values, shown by an example 4-somite-pair embryo. C: (i) PCA dimension reduction of the smoothed TF values in order to identify cell states. PCA loadings per TF indicated by arrow direction and magnitude (n=19 embryos). (ii) The pseudotime tool Slingshot (Street et al., 2018) identifies a path between the SOX2 high and the TBX6 high regions to create a pseudo-space continuum (n=19 embryos). D:) The pseudo-space continuum values (corresponding to cell states as defined by expression of TBXT, SOX2 and TBX6) mapped back onto the manifold projection of the four-somite-pair epiblast. Pseudo-space values within known N-M-bifated regions are likely to correspond to NMP identities. E: Red nuclei correspond to cells with pseudospace values defined in (e) as corresponding to likely NMP identities (‘NMP-like cells’). These map to a U-shaped region that extends beyond the boundaries of the known N-M-bifated regions.

The NMP region can also be defined based on co-expression of SOX2 and TBXT [32,42]. However, this marker-based approach presents challenges because both TBXT and SOX2 are expressed in graded distributions, meaning that gating for positive cells can be somewhat arbitrary. For example, our manual gating of TBXT+SOX2+ double positive cells in E8.5 embryos (brown cells in Figure 3A) defines a region of double positive cells that extends some distance beyond the known N-M-potent regions (Figure 3A). Furthermore, this analysis produces ill-defined borders between neighbouring regions due to local heterogeneity in expression of TBXT and SOX2 (Figure 3A). We therefore sought an approach to harmonise the known N-M-potent regions with the SOX2/TBXT/TBX6 expression profile in the epiblast. Our aim was not to definitively assign NMP character to each individual cell, but rather to approximate a discrete region that is likely to contain cells of NMP identity.

We first quantified TBXT, SOX2 and TBX6 fluorescence immunofluorescence intensity for each nucleus in our segmented data (described in Figure 2). Then, values obtained from different images were normalised to enable cross-image comparison and smoothed using a moving average across local neighbourhoods to reduce local noise (Figure 3B, Methods). Then, normalised and smoothed values of TBXT, SOX2, and TBX6 expression from multiple embryos were subjected to PCA analysis (Figure 3Ci). The pseudotime tool slingshot [43] was used to isolate a trajectory from SOX2+ to TBXT+SOX2+ to TBXT+TBX6+ to align nuclei along a TF pseudospace axis (Figure 3Cii), representing a continuum of cell states. Our aim here was not to define a temporal differentiation trajectory but rather to generate a range of values representing different cell pseudospace identities. These values were then mapped back onto the “PRINGLE projection” epiblast (Figure 3D) in order to establish which pseudospace identities correspond to the regions of known NMP potency (black boxes in Fig 3D). On this basis, these values were assumed to represent an NMP-like identity.

Nuclei that were defined in this way as having NMP-like identity mapped to the epiblast projection in a U shape that extends a short way beyond the known NMP regions, potentially encroaching on the neural-fated lateral border (Fig 3E, Supp Fig 3). This approach provides a relatively unbiased method to define regions of interest based on combining information from multiple graded markers and relating this information to regions of experimentally-determined fate. Comparison of the fate map with the pseudospace map allowed us to approximate a region in pseudospace that approximates to the NMP-containing region, providing a framework for defining patterning events in relation to this region.

### Mapping the shape and steepness of gradients of transcription factor expression in 3D

We next asked how patterning of SOX2, TBXT and TBX6 relates to the position of the putative NMP region. This information is useful because, for example, steep changes in TF expression in time or space could indicate when and where cells are likely to be making differentiation decisions. To measure graded changes in marker expression at high spatial resolution, we compared TF expression values between neighbouring cells (Figure 4A: see Supp Figure 4 and supplemental methods for a full description of this approach).

**Figure 4.**
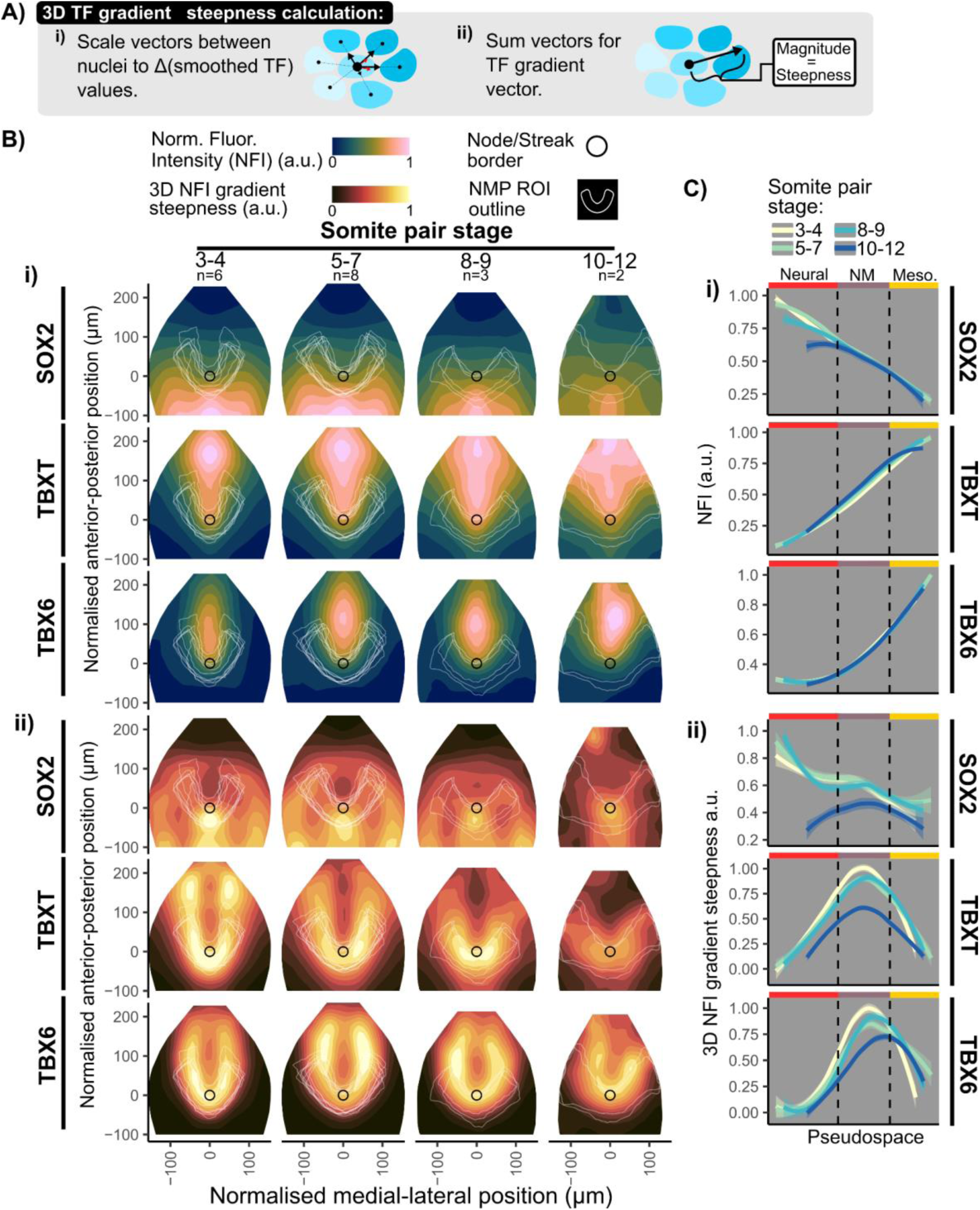
Mapping the shape and steepness of gradients of transcription factor expression in 3D. A:. Method to map the shape of gradients across 3D space (i) First scale the unit vectors between a nuclei and its neighbours to the change in smoothed TF values. (ii) Then sum the vectors to obtain a single directional vector, with magnitude representing the steepness of the gradient. B: (i) Contour plots of average binned normalised fluorescence intensity (NFI) measurements for SOX2, TBXT and TBX6 in the normalised epiblast shape using PRINGLE, as described in Figure 2. Embryos are grouped per somite pair stage during E8.5, n numbers shown per group. White lines indicate the NMP region of interest (ROI) for each individual embryo as determined using the approach shown in Figure 3. (ii) Contour plots mapping TF gradient steepness map, calculated as shown in A. Averaged data from multiple embryos is shown, as explained in (c). Gradients of TBXT and TBX6 appear to be steepest in regions corresponding to NMP-like cells (NMP ROI) and gradients of SOX2 appear to be steepest in the node streak border. C: i) Averages of single cell NFI measurements, as measured in Figure 2a, and measurement of gradient steepness, as measured in b (ii), per embryo and somite pair stage plotted in relation to pseudo space values, corresponding to cell identities based on integration of SOX2, TBX6 and TBXT expression as determined in Figure 3. This confirms that gradients of TBX6 and TBXT gradient are steepest within the NMP ROI and decreases into the PS, with TBX6 spatially lagging behind TBXT. Solid lines indicate the fit of a non-parametric multiple regression curve and shading indicates 95% confidence intervals. Vertical dashed lines indicate gating from fitting TF pseudospace to the N-M-bi-fated regions in Figure 3.

We first used the methods described in Figure 2 to computationally flatten and align epiblasts to make it possible to display averaged patterns across multiple embryos, visualised as manifold projections of the epiblast (PRINGLE projections). We then used the method described in Figure 3 to establish the putative NMP-containing region for each individual embryo (white U-shaped outlines in Figure 4B & Figure S3). Embryos were staged according to the number of somite pairs (SP), and the analysis described below is presented as the averaged pattern across multiple stage-matched embryos from 3 to 12 somite pair stages. (Figure 4B).

Mapping the fluorescence intensity values onto averaged epiblast projections revealed the complex shapes of the gradients of SOX2, TBXT and TBX6 expression over time. SOX2 exhibits an M shaped distribution, extending caudolaterally into the caudal lateral epiblast. Its expression shifts anterior-wards relative to the node between 4SP and 10SP stages. TBXT is highest at the posterior midline, decreasing anteriorly and laterally into the caudal lateral epiblast and anterior streak to form a V shape; this domain expands over time relative to the size of the epiblast, with the sharpness of the V shape flattening out somewhat at 10-12 SPs. TBX6 forms an elliptical expression domain centred on the middle of the primitive streak (Figure 4Bi)

We then asked where each marker exhibits the steepest change in expression. To quantify this, we calculated the magnitude of changes in smoothed fluorescence intensity values between a cell and its neighbours in 3D. (Figure 3c ii, Supp Figure S4). The resulting values were projected onto the manifold projection of epiblasts in order to visually highlight the regions with steepest changes in fluorescence intensity (Fig 4Bii). This analysis reveals that TBX6 expression changes sharply around the medial regions of the NMP region at all stages examined. TBXT expression also changes sharply in a similar region to TBX6 at SP 4,6 but expression flattens out somewhat by SP10-12. In contrast, SOX2 expression changes most prominently at the lateral and anterior edges of the NMP region (Figure 4Bii).

Finally, we used our pseudospace measure of state (Figure 3) to ask how fluorescence intensity values and local gradient steepness for each of the three transcription factors relate to cell identity (Figure 4C). In summary, we were able to use an approach based on local-neighbour-comparisons to map the steepest changes in SOX2 expression to a region where cells are committing to a neural fate, and to identify sharp changes in TBXT and TBX6 in regions where cells begin to adopt a mesoderm fate (Cambray and Wilson 2007, Wymeersch et al. 2016).

### Mapping the properties of cells in relation to their local neighbourhood

Having explored graded changes of TF expression, we next examined mesoscale heterogeneity. In particular, we asked when and where cells tend to be different from their immediate neighbours (referred to here as “local variability”). This information is useful because, for example, the emergence of isolated TBX6+ cells within the epiblast could be consistent with sporadic mesoderm commitment in some cells followed by lateral inhibition of differentiation in neighbours.

We used the analysis described in Figure 2 (without smoothing between neighbours) to establish fluorescence intensity values for SOX2, TBXT and TBX6 in each nucleus and to identify the nearest neighbours for each nucleus in 3D. We then used the coefficient of variation (CV) of TF values between a cell and its neighbours in order to obtain a simple relative measure of how similar each cell is to its local neighbourhood (Figure 5A). This revealed that TBX6 exhibits the most local spatial heterogeneity, and SOX2 the least (Figure 5B).

**Figure 5:**
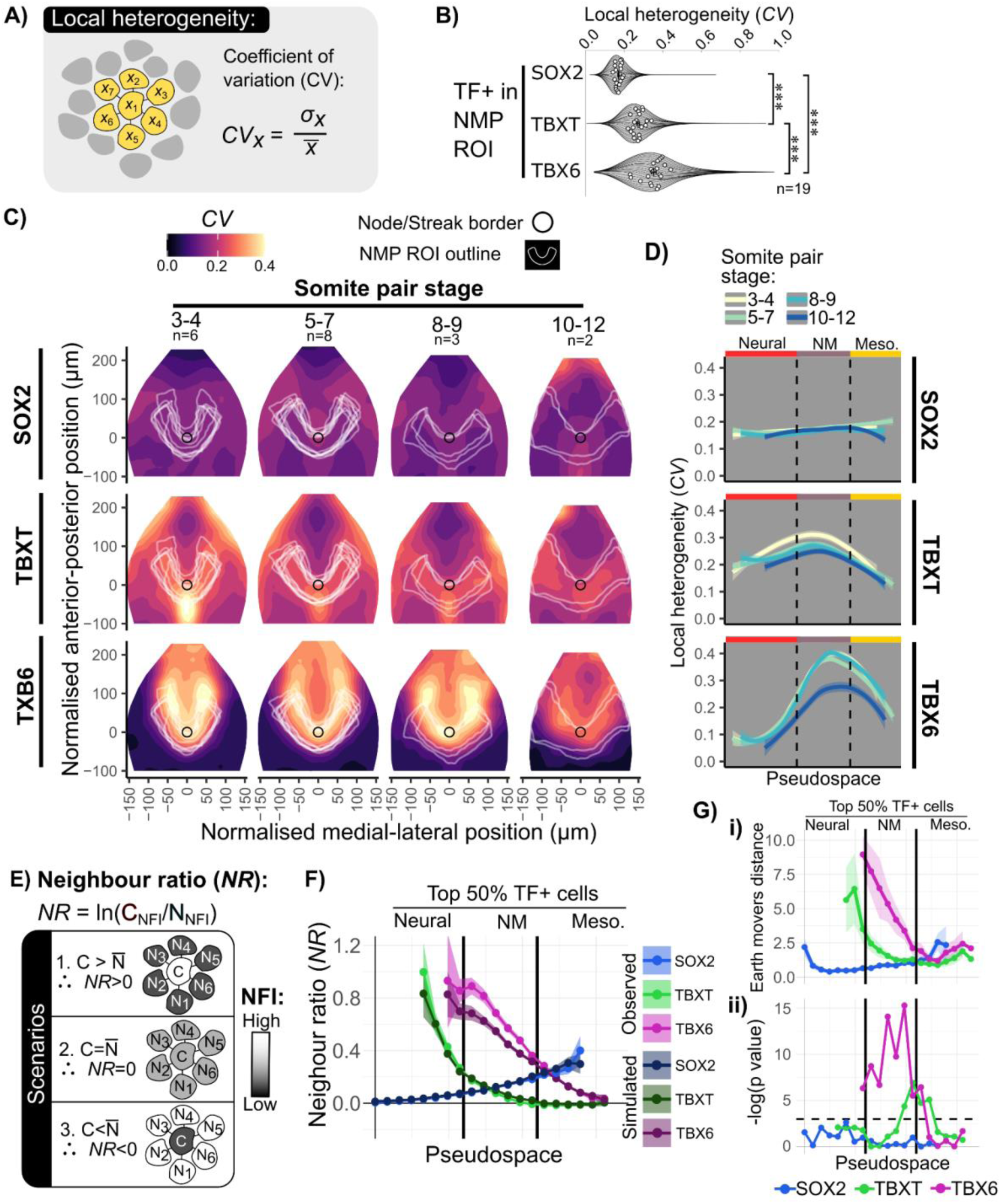
Mapping the properties of cells in relation to their local neighbourhood **A:** The coefficient of variation (CV) is used as a metric for the degree of local heterogeneity in fluorescence intensity values for a cell and its direct neighbours (in 3D), defined as the standard deviation divided by the mean of fluorescence intensity values for a cell and its neighbours. Neighbours are defined as in Figure 2A. **B.** Violin Superplots showing CV values for SOX2, TBXT and TBX6 within NMP-like cells as defined in Figure 3. Local heterogeneity (CV) in TBX6 is greater than TBXT, which is greater than local heterogeneity in SOX2. Dots indicate the mean for each of 19 embryos. Statistical tests performed with Tukey’s HSD. *** = p<0.001. **C:** Maps of local variability in TF expression across the epiblast. Contour plots showing CV values mapped onto EpiMap. Data is averaged from multiple embryos as described in Figure 4). U-shaped white lines indicate the NMP ROI as measured in Figure 3. High local variability (CV) for TBXT and TBX6 is observed within the NMP ROI. **D:** Heightened TBXT and TBX6 CV to the NMP region and decreases into the mesodermal PS. Solid lines indicate median and dashed lines indicate 95^th^ percentile per embryo. **E:** The neighbour ratio (NR) metric can distinguish patterning scenarios where a cell is (1.) surrounded by cells with an averaged expression that is higher expression than itself, (2) the same as itself, or (3) lower expression than itself. Defined as ln (Cell TF value/ Average Neighbour TF value). **F:** Comparing the ‘high’ cells in the TF+ populations (upper 50th percentile) for SOX2,TBXT and TBX6 to synthetic data assuming a normal distribution for a cell’s expression and its neighbours. This shows a decrease in NR, indicating TBX6+ high cells are surrounded by cells expressing lower TBX6 levels than expected within a random pattern. Points indicate average NR per embryo, shaded areas show confidence intervals of 0.05 and 0.95. **G:** Statistical analysis of the difference in observed and synthetic data NR values along pseudospace bins using (i) earth mover’s distance between normalised distributions for each embryo and (ii) p values resulting from one way ANOVA between observed and synthetic embryo means. The difference in TBX6 NR values is statistically significant in the NMP ROI with a high difference in distributions compared to TBXT or SOX2. Shaded areas show confidence intervals of 0.05 and 0.95. Dashed line in ii) indicates p-value = 0.05.

We next asked whether high CV values map to particular embryonic regions. We used methods described in Figure 2 to computationally flatten and align epiblasts in order to display averaged patterns across multiple embryos, visualised as manifold projections of epiblasts. We then used the method described in Figure 3 to establish putative NMP regions for each individual embryo (white U-shaped outlines in Figure 5C). We then mapped CV values (Figure 5A, B) onto epiblast projections (Figure 5C). Embryos were staged according to the number of somites, and analysis is presented as the averaged pattern across multiple stage-matched embryos from 3 to 12 somite pair (SP) stages. (Figure 5C: note that these are the same embryos that are analysed in Figure 4).

This analysis confirmed that SOX2 exhibits low CV values across the region examined, while TBXT exhibits modest CV values within the NMP region and a hotspot of local heterogeneity around the anterior tip of the primitive streak. In contrast TBX6 has high CV values mapping to the medial edges of the putative NMP region (Figure 5C). Plotting CV values for each cell in relation to pseudospace measure of cell identity (Figure 5D) confirms that local variability in TBX6 is associated primarily with the putative NMP region.

We then sought to distinguish between three patterning scenarios: a cell is surrounded by (1) higher, (2) the same, or (3) lower TF values compared to itself (Figure 5E). These scenarios can be distinguished by measuring the ratio between fluorescence intensity of each cell to the averaged fluorescence intensities of its neighbours (Neighbour Ratio: NR). In order to facilitate comparison between different cells, the log of NR is calculated to ensure proportional influence on NR for changes in numerator (cell) and denominator (average neighbour) (Supp Figure S5). To evaluate whether the observed NR values are consistent with a random distribution, synthetic data was produced assuming a normal distribution of TF values between a cell and its neighbours (Supp Figure S6). For example, observed and synthetic values for the top 50% TBX6+ cells were plotted for each cell in relation to their Pseudospace identity as determined in Figure 3 (Figure 5F: see Figure S6 for more details). We then computed the earth move distance between the observed and synthetic NR distribution to obtain an informative measure of differences between these distributions (Figure 5G)

This analysis (Figure 5F) indicates that, within the putative NMP region, SOX2 and TBXT have relatively low NR values and that their local neighbour relationships are consistent with a random distribution. In contrast, within the putative NMP region, this analysis indicates a non-random association of TBX6-positive cells with TBX6-negative neighbours.

This analysis method offers a quantitative approach to assess and compare local coherence vs heterogeneity in patterning of cell states. It confirms that SOX2+ and TBXT+ cells tend to be located in relatively coherent neighbourhoods, i.e. they are mostly surrounded by cells of similar identity. In contrast, TBX6 exhibits considerable local heterogeneity within medial regions of the putative NMP domain, with isolated TBX6-high cells surrounded by TBX6-low cells. This raises the possibility that TBX6 expression is regulated by intercellular feedback signalling that enforces local cell fate diversification.

### Measuring local heterogeneity within embryos, gastruloids and monolayer differentiation

ES-cell-derived models of development are tractable experimental systems for studying cell-fate diversification but may not recapitulate all aspects of embryonic patterning. Having found that TBX6+ high tend to be surrounded by TBX6-low cells within the posterior epiblast of E8 mouse embryos (Figure 5), we asked to what extent this pattern is recapitulated in two commonly used in-vitro model systems: 2D monolayer differentiation of hES cells into NMPS [44,45] (Figure 6A), and 3D differentiation of mESCs into gastruloids [46,47] (Figure 6B).

**Figure 6.**
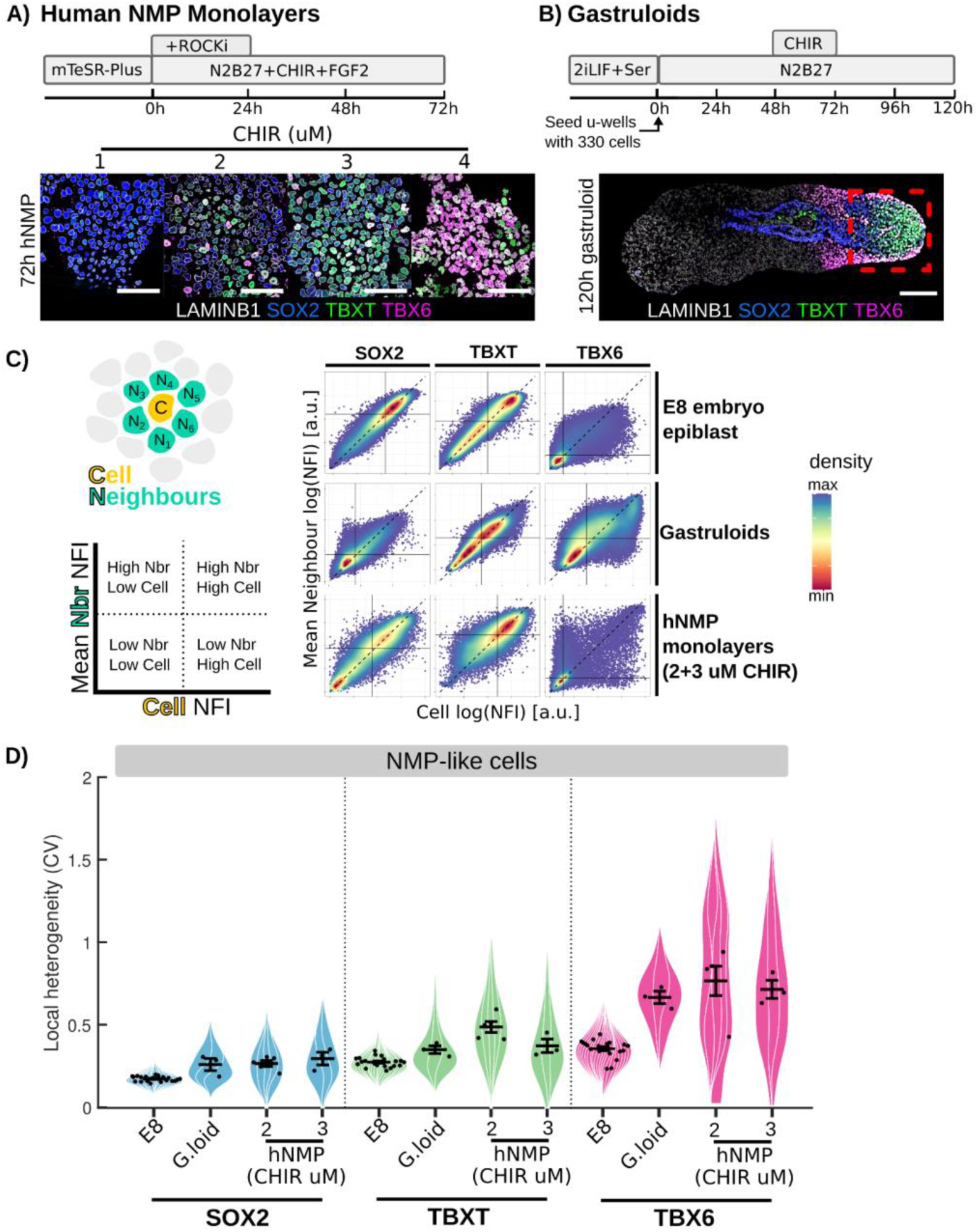
Comparing local patterning between embryos, gastruloids and monolayer cell culture. A: Protocol for generating hNMPs from hESC and representative confocal images of the culture fixed at 72h and stained for SOX2, TBXT, TBX6 and LAMINB1 (further characterisation fig SX). B: Protocol for generating gastruloids [47] with a representative confocal image of a gastruloid fixed at 120h for SOX2, TBXT, TBX6 and LAMINB1. Red box indicates typical field of view for high resolution image acquisition and subsequent neighbour analysis. C: Density plots comparing each nucleus’s TF normalised fluorescence intensity (NFI) value to its neighbours shows strong linear correlations for TBXT and SOX2, but not for TBX6 with many ‘high’ TBX6 cells surrounded by ‘low’ TBX6 cells. Consistent in the E8 epiblast, gastruloids, and in vitro human NMP like cell monolayers. Black line indicates the median and the dashed line indicates x=y line. D: Violin superplots of TF local heterogeneity (coefficient of variation [CF]) for respective TF+ cells within NMP-like cells isolated in gastruloids (process outlined in Fig S8) and hNMP monolayers compared to in vivo WT E8 bi-fated region. NMPLC populations within in vitro models are more heterogeneous than comparative in vivo NMP populations. All scale bars indicate 50µm. Points represent replicate means. Error bars represent 5% to 95% confidence intervals. Gastruloids n= 3 with 3 technical replicates.

Comparing patterning of embryos with patterning of cell-culture-based systems is challenging due to difficulties in identifying the particular populations in culture that correspond to a region of interest in vivo. For embryos, we used spatial cues to define the posterior epiblast and then defined the NMP region by combining information from marker expression with knowledge about potency based on grafting experiments (as described in Figure 2). For monolayer differentiation cultures, we used Chiron to impose an NMP character (Supplemental Figure S7) and consider the entire culture to be our region of interest. For gastruloids, we focused on the posterior region of the structure (red box in Figure 6B) and then identified NMP-like cells using an approach similar to that outlined in Figure 2, although in this case we relied exclusively on marker expression (Supplemental Figure S8) because the potency of particular subregions within gastruloids has not been defined.

We confirmed that NMP differentiation from hESC was efficient at concentrations of 2 or 3uM CHIR99021 (Chiron) based on co-expression of TBXT and SOX2. (Supplemental Figure S7). We also confirmed successful emergence of NMP-like cells within the tip of gastruloids generated from mESC (Supplemental Fig S8). Interestingly, the NMP-like cells most resembling those defined in vivo (Figure 3B-D) are located proximally to mesoderm cells at the tip of the elongated gastruloid (Supplemental Fig S8), reminiscent of the location of NMPs within the tail bud [41]. We then compared local variability of SOX2, TBXT, and TBX6 within both these in vitro systems to data obtained from the epiblasts of E8 embryos (embryo data was obtained as shown in Figure 5).

We calculated normalised fluorescence intensity (NFI) values for SOX2, TBXT and TBX6 in each nucleus and identified nearest neighbours in gastruloids and monolayer cultures using the same pipeline as for embryos (see Methods and Fig 2A). We then plotted the NFI for each cell against the mean NFI of all its neighbours in order to visualise how similar each cell is to its local neighbourhood (Figure 6C). This analysis (Figure 6C) confirms that, of the three TFs examined, TBX6 exhibits the most local spatial heterogeneity (many points do not align along the diagonal), and SOX2 the least spatial heterogeneity (most points align along the diagonal) in all three experimental systems. In the case of embryo and gastruloids, more cells lay beneath the diagonal than above it, indicating that TBX6-Hi cells tend to be surrounded by TBX6-low cells in both cases. This pattern was not recapitulated for monolayer differentiation cultures. To more carefully compare heterogeneity between different systems, we calculated the coefficient of variation (CV) of TF values between each cell and its neighbours (see Figure 5A). We then plotted the mean CV for each culture, gastruloid, or embryo (Figure 6D). This analysis confirms that hES-derived monolayer differentiation in unconstrained cultures exhibit a higher degree of local variability in TBX6 when compared with embryos or gastruloids. TBX6 heterogeneity in gastruloids was lower than in monolayer cultures but higher than in embryos.

We conclude that our analysis methods are useful for comparing local variability between different experimental models, although a detailed like-for-like comparison would require knowledge of the fates of cell neighbourhoods that is currently not available in cell culture model systems.

### Local heterogeneity in TBX6 is sensitive to Notch inhibition

One advantage of our analysis pipeline is that it allows quantitative assessment of changes in patterning after experimental manipulation of candidate regulators. It has recently been reported that Notch signalling can influence differentiation of hES-derived NMPs [39]. Notch mediates lateral inhibition to generate heterogeneity in many other contexts [10,39], making it an excellent candidate for contributing to the heterogeneity in TBX6 that we observe within the NMP region (Figure 5). We therefore asked if we could use our image analysis pipeline to detect changes in TBX6 patterning after pharmacological manipulation of Notch activity.

Posterior explants of E8.5 embryos were cultured with Notch signalling inhibitor LY or with vehicle-only (DMSO) for 12 hours (Figure 7A). Fluorescence intensities were measured for SOX2, TBXT and TBX6. Embryos were analysed using the approaches described in Figures 2-5 (Figure 7A,B). We confirmed that embryo explants displayed broadly normal tissue morphology, similar to a late E8.5 or early E9.0 embryo, after the short 12h culture period (Figure 7B), indicating that the overall shape and size of the epiblast region was not grossly affected by Notch inhibition.

**Figure 7.**
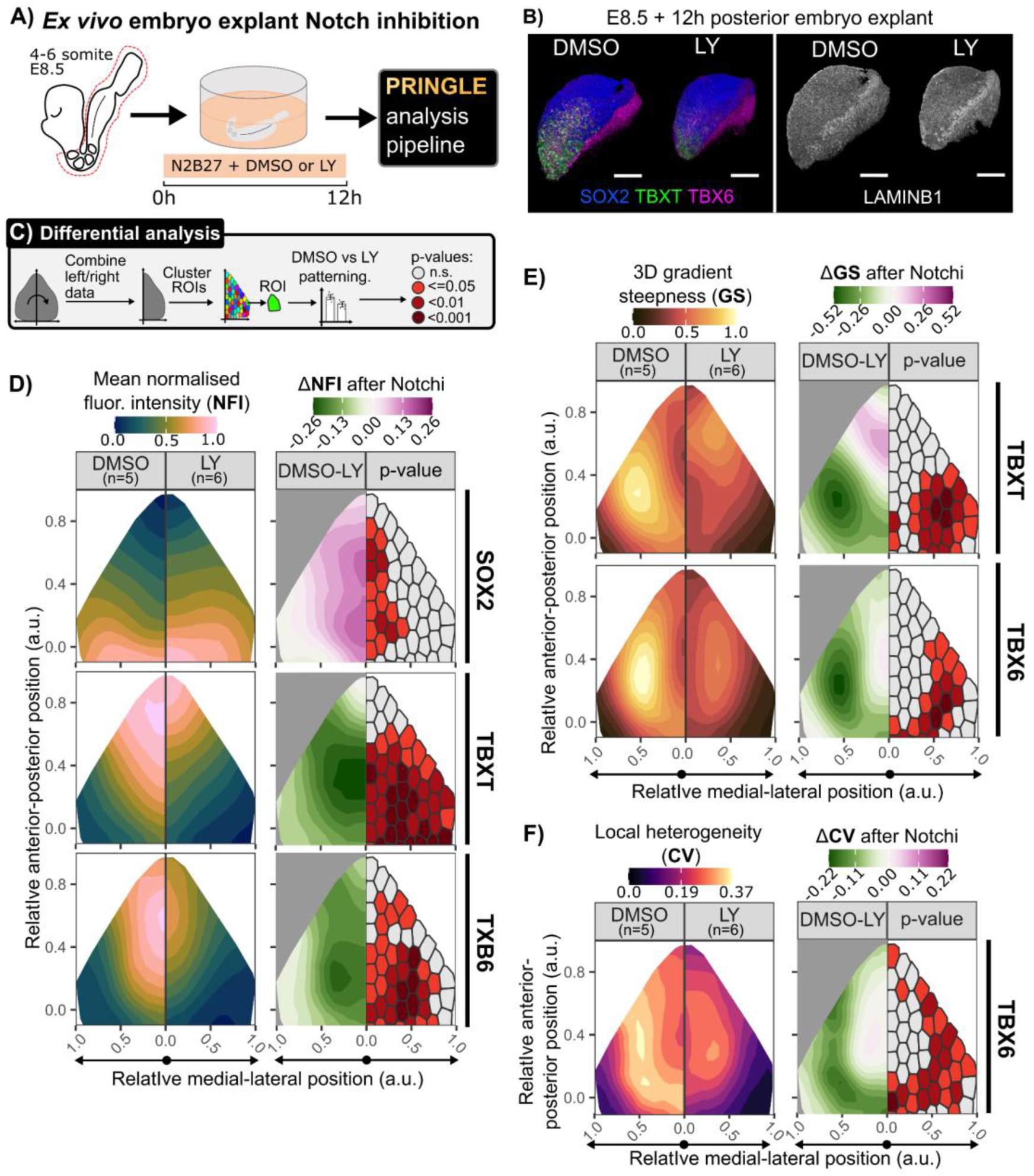
Mapping changes in patterning after inhibition of Notch activity. A: Culture method where a posterior explant of an E8.5 embryo is cultured in N2B27 media with the Notch inhibitor LY at 150nM or an equal amount of DMSO for 12 hours. B: Representative 3D renders of explants for each condition, stained for LaminB1 or SOX2, TBXT, and TBX6 indicating normal tissue morphology but a reduction in TBXT and TBX6 expression. Anterior (A), posterior (P), dorsal (D), ventral (V) axes are indicated. Scale bars indicate 100μm. C: Analysis pipeline to evaluate the effect of Notch inhibition on patterning. After using PRINGLE to create an EpiMap, the epiblasts are folded at the midline and clustered with k-means to demark comparable ROIs. Patterning metrics are compared between DMSO and LY in these regions. One way ANOVA is conducted to identify the statistical significance of the difference of patterning metrics, regions are filled with associated colours to denote the p-value. D: Contour plots of TF normalised fluorescence intensity (NFI) projected onto the EpiMap with the difference in DMSO vs LY conditions and associated p-values per ROI. Displaying an increase of SOX2 NFI in the PS and a posterior withdrawal of the TBXT and TBX6 domains in the Notch inhibited condition. E: Contour plots of NFI gradient steepness projected onto the EpiMap with the difference in DMSO vs LY conditions and associated p-values per ROI. Note the overall decrease in TBXT and TBX6 gradient steepness, with a new posteriorly located nonlinear TBXT gradient. F: Contour plot of TBX6 local heterogeneity (CV) on EpiMap. Note the statistically significant decrease in the CLE and NSB like regions for both TBXT and TBX6.

To visually and quantitatively compare control (DMSO) and Notch-inhibitor-treated (LY) epiblasts, we used the following procedure: First, we took advantage of the bilateral symmetry of the embryo and combined the left and right portions of our PRINGLE projections into one single projection (Figure 7C). This allowed us to provide a compact view of our data for side-by-side comparison between treated and control conditions (Figure 7D-F).

Next, we used k-means clustering to partition projections into bins of approximately 100 cells. This binning approach allowed us to compute the statistical significance of differences observed between conditions and report the spatial distribution of p-values on the PRINGLE projection.

Control DMSO-treated embryos show a distribution of SOX2, TBXT and TBX6 similar to that previously observed in 8-10SP E8.5 embryos (compare Figure 4Bi with Figure 7D). In contrast, after Notch inhibitor treatment, the SOX2 expression domain slightly expanded anteriorly while expression domains of T and TBX6 were reduced, with the most significant difference observed within lateral epiblast regions that would normally be expected to contain NMPs (see Figure 3). This is broadly consistent with a role for Notch in promoting mesoderm fate at the expense of neural differentiation [39].

We then measured the shape and steepness of TF gradients, as described in Figure 4. Notch inhibition brought about a significant decrease in the steepness of TBX6 and TBXT gradients within the more anterior regions of the CLE, consistent with a depletion in NMPs overall. Finally, we measured local heterogeneity in TBX6 as described in Figure 5. Strikingly, although TBX6 was still expressed in the presence of Notch inhibitors, local heterogeneity was greatly reduced.

We conclude that patterning of NMPs and their early mesoderm derivatives is sensitive to Notch inhibition. This showcases the utility of our patterning analysis as a useful means to determine the effect of developmental regulators on aspects of patterning, at tissue-scale and at the level of individual cells and their neighbours.

## Discussion

Quantification of local patterning within tissues is a non-trivial task, even when analysis is restricted to 2D images [48]. Extending analysis to three dimensions presents additional challenges. Here by measuring how similar each cell is to its local neighbourhood in 3D, we could determine the local direction and steepness of graded transcription factor expression and extract quantitative descriptions of fine-grained heterogeneity in cell identity. We used these methods to examine when, where, and how changes in cell identity occur in the posterior of mouse embryos at the onset of axis elongation.

Our image analysis pipeline allows spatial comparison between samples at single cell level. It is often challenging to find robust ways to compare image data. Standardised micropatterns offer one solution to quantify reproducibility of patterning and to reliably measure the effects of experimental manipulations in experiments using cultured cell monolayers [49]. It is considerably more challenging to quantitatively compare patterning between embryos or 3D cultures because these have complex and somewhat variable shapes and sizes: for example, the E8.5 epiblast of the mouse embryo has a complex curved shape. This problem can be addressed with methods to project complex curved shapes onto 2D planes [50]. Here we developed a simpler approach that is based on analysis of segmentation data rather than raw images, and which incorporates a module to normalise the spatial data to embryonic axes. This establishes a framework within which to quantify patterning across multiple embryos.

In order to compare local patterning between embryos, it is first necessary to define a region of interest within which patterns are to be compared. However, regions of interest are often not clearly demarcated by coherent expression domains with distinct boundaries. One such example is the region of the embryo that harbours neuromesodermal progenitors during axis elongation. In this study, we develop a method to integrate information from multiple graded markers with previous experimental data defining the fate of microdissected regions [40,41]. This establishes T/Sox levels that define putative NMPs. Our approach here builds on a previous study [41] which used a simpler thresholding approach to define a U shape describing mid-level T and SOX2 co expression at E8.5. Here we advance on that approach by avoiding arbitrary thresholding of individual markers.

The putative NMP domain we identify includes part of a region lateral to the node-streak border previously assumed to contain neural-committed progenitors. This may explain why this region shows two alternative contribution patterns: one, where cells exit en masse to the neural tube, and the other, where cells are retained in the tail bud and contribute to neurectoderm and mesoderm: the dissected region contains cells of two alternate phenotypes: High SOX2/negligible TBXT neural-committed cells towards the anterior/lateral edge of the rectangular shape dissected for grafting, and mid SOX2/TBXT NMPs at the posterior/medial side. The grafting procedure partially disperses cells in these regions, which may have resulted in some grafts retaining NMPs while others contain only neural-committed cells. Significantly, we also find that coexpression of high TBXT/low SOX2 at the midline occurs in a mesoderm-committed region, suggesting that mesoderm commitment is associated with a threshold level of TBXT rather than extinction of SOX2.

Interestingly, the gradients of all three TFs are steepest at the anteriormost part of the NMP region near the node. This locale contains the longest-lived NMPs (Cambray and Wilson 2007), and contacts the most posterior notochord progenitors ventrally. Heterotopic transplants of NMPs show that contact with the posterior notochord progenitors increases the retention of NMPs as progenitors in the tailbud, without compromising their ability to differentiate (Wymeersch et al eLife 2016). Furthermore, the posteriormost notochord progenitors are vital for axial elongation [51]. The presence of steep gradients of all three TFs in this location is consistent with an NMP population anchored via a small and stable niche, the posterior notochord, from which cells can readily exit towards both neurectoderm and mesoderm.

The mechanisms that regulate mesoderm commitment are not fully understood. Mesoderm differentiation is driven by Wnt and FGF signalling, yet these pathways appear to be active throughout the caudal epiblast [52–55], with no evidence for upregulation of activity at the primitive streak. Furthermore, Wnt and FGF are required for maintenance of progenitors as well as their differentiation into mesoderm [32,33]. It therefore seems likely that additional factors must influence mesoderm commitment. Notch is one promising candidate: Notch ligands and targets are upregulated at the midline, and Notch can regulate mesoderm commitments [33,39,56].

In keeping with the idea that Notch regulates mesoderm commitment, our neighbour-comparisons reveal that mesoderm-commitment marker TBX6 is expressed in a spotty pattern within the NMP region. Isolated TBX6+ cells are non-randomly surrounded by TBX6-neighbours, and this pattern is sensitive to pharmacological suppression of Notch activity.

Based on these findings an attractive hypothesis is that lateral inhibition through Notch [10] balances commitment to mesoderm fate with preservation of a progenitor pool as cells approach the midline. However, we cannot exclude other mechanisms: local variability between neighbours does not necessarily indicate lateral inhibition of cell identity: it could represent bistability in stochastic cell fate allocation at threshold levels of signalling inputs [57], or it could be the consequence of migration of cells from an initially coherent domain [58]. It would be interesting to use emerging neighbour-labelling technologies [59–61] to test more directly whether, for example, TBX6+ cells suppress mesoderm commitment in their neighbours.

In summary, our patterning-analysis methods can be used to gain new insights into regulation of cell identity during early post-gastrulation mouse development. These approaches should be readily applicable to any system that is amenable to single cell segmentation, and it will be particularly interesting to extend these approaches to single-cell-resolution spatial transcriptomics datasets. We present these approaches as a toolkit for studying patterning during development, homeostasis, regeneration or disease.

## Acknowledgements

We are grateful to all members of the Lowell, Wilson and Blin labs and to Linus Schumacher for helpful suggestions. We thank facility staff at the Institute for Regeneration and Repair of the University of Edinburgh, especially, Justyna Cholewa-Waclaw at the High-Content Screening Facility, Matthieu Vermeren at the Imaging Facility, and Theresa O’Connor at the Tissue Culture Facility. MF was supported by an EastBio PhD studentship; SL is supported by a Wellcome Trust Senior Fellowship [220298]; VW is supported by MRC grant MR/S008799/1; GB is supported by a BBSRC grant ref:BB/W002310/1, and a WT ISSF3 award ref: IS3-R1.16 19/20; JKD is supported by MRC grant MR/X018423/1

## Methods

### Embryo dissection and culture

Mice were bred and housed in the Animal Unit of the Centre for Regenerative Medicine in accordance with the provisions of the Animals Act 1986 (Scientific procedures). Pregnant female mice were culled by cervical dislocation by the Animal Unit staff. The uterus was dissected from the mice and placed into M2 media at RT, whereafter the decidua and Reicherts membrane were removed using forceps.

For embryo culture, the anterior portion up to the first somite was removed while in M2 media. These were immediately transferred to N2B27 media supplemented with Pen/Strep antibiotics and either 150nM LY411575 or an equal amount of DMSO. These were cultured statically in an incubator at 37°C and 5% CO2 for 12 hours.

### Mouse ES and gastruloid culture

E14Ju09 ES cells [62] were routinely maintained in FCS (Gibco) with 100u/ml LIF (produced in-house) on gelatinised plates as described in [59]. Prior to gastruloid formation, ES cell culture media was additionally supplemented with 3μM CHIR 99021 and 5μM PD 0325901 5μl 1μM (2i LIF FCS media) for at least 10 days in order to reduce spontaneous differentiation.

Gastruloids were generated as previously described [47]. 2i/LIF/FCS ES cells are washed in PBS before adding 0.05% Trypsin EDTA solution. After ES colony detachment, 5-10 volumes of fresh 2i/LIF/FCS were added to quench the Trypsin EDTA solution. The cells and media were transferred to a universal tube and pelleted by centrifugation at 300g for 3 minutes. The media was aspirated, and the pellet was resuspended in cold PBS, making sure to create a single cell suspension. This PBS wash was repeated once more, then the cell pellet was resuspended in prewarmed N2B27 to create a single cell solution. The cells were then counted and then a solution of N2B27 medium with 8,250 cells/mL was made. 40uL of this medium, containing ∼330 cells, was plated into untreated u-bottom 96 wells and incubated for 48 hours at 36 °C and 5% CO2. At 48 hrs, 150uL of N2B27 and 3uM CHIR99021 (Axon) were added into the well, the last half of the media was expelled forcefully to dislodge the aggregate but without spilling the medium. The aggregates were cultured for a further 24 hrs to 72 hrs, after which 150uL of media was removed and replaced with 150uL fresh N2B27. From 96h to 120h the media was replaced with fresh N2B27 supplemented with DAPT 50μM or an equal amount of DMSO. At 120h the gastruloids in each condition were fixed.

### Human ESC culture and hNMP differentiation

The clinical grade hESC cell line MasterShef7 was used (Vitillo et al., 2020). MasterShef7 hESCs were cultured on Geltrex™ (Gibco) coated wells in mTeSR Plus™ media. Routine hESCs passaging was performed with accutase in a 1:10 ratio once 70-80% hESC confluency was reached, typically every 4-5 days. mTeSR Plus™ media was supplemented with ROCK inhibitor Y-27632 10μM for the first 24 hours after passaging. hESCs were differentiated to hNMP-LCs as described in [44,45].. hESCs were plated at 10,000 cells/cm2 on Geltrex™ in N2B27 media supplemented with Y-27632 10uM with the desired concentration of recombinant FGF basic (R&D systems) and CHIR99021 (Axon). Typically, 20ng/mL FGF and 2uM or 3uM CHIR99021 was used. After 24 hours, the media was aspirated and fresh prewarmed N2B27/FGF/CHIR media without Y-27632 was added. hESCs were cultured for a further 48 hrs to acquire hNMP-LCs identity at 72 hrs.

### Immunohistochemistry

For E8.5 embryos and cultured embryos, the posterior portion from the posteriormost somite to the allantois were sub-dissected immediately prior to fixation. Embryo posterior portions and gastruloids were briefly washed in PBS and then fixed in 4% PFA in PBS + 0.1% Triton for 2 hrs at room temperature with gentle agitation. hNMP-LC monolayers were fixed by adding 8% PFA PBST to an equal volume of media retained in culture for a final 4% PFA concentration for 15 minutes. All tissues were washed in PBST and permeabilised in 0.5% Triton + PBS for 15 minutes, then placed in blocking solution at 4°C overnight. All tissues were costained for SOX2 (Abcam, Ab92494), TBX6 (R&D Systems, AF4744), T-NL557 conjugate (R&D Systems, NL2085R), and LaminB1 (Abcam, Ab16048) conjugated to Alexa-Fluor647 using a conjugation kit per the instruction manual (Thermofisher, A20186). Sequential stains were performed by first staining SOX2 + TBX6 primaries, followed by donkey anti Rabbit Alexa Fluor 488 (Abcam, A21206) and donkey anti goat Alexa-Fluor594 (Invitrogen A21206) secondary for 24 hours with gentle agitation. The secondary was blocked with 5% Rabbit + 10% Goat + 5% Donkey serum in PBST (PBST-RGD). The second primary stain with conjugated T-557 NL and LaminB1-647 were performed in PBST-RGD blocking solution. All primary and secondary stains were performed at room temperature with gentle agitation. 3D tissues were stained for 24 hours and monolayers for at least 4 hrs. Three 15-minute washing steps in PBST were performed at RT with gentle agitation between each step.

For hNMP high throughput single cell quantification, cells were routinely passaged and replated at half density onto Geltrex coated wells with media composition the same as the pre-passage culture media. Cells were left to adhere for 10-15 minutes and then immediately fixed as described above. The cells were stained with DAPI (Sigma, D9542) diluted 1:10,000 in blocking solution. Cells were stained as described above, except TBX6 secondary staining is performed with Donkey anti-goat Alexa-Fluor647 (Invitrogen, A21447).

Confocal microscopy was performed after tissue dehydration in a PBS/methanol series (5 minutes each) and two 10 min 100% ethanol steps. Dehydrated tissues were transferred to BABB (2:1 benzyl alcohol:benzyl benzoate) in ibidi µ-Slide wells for imaging. Monolayer tissues were mounted in 90% glycerol 10% PBST for confocal imaging.

### Image analysis

#### 3D Nuclei segmentation and curation

3D single nuclei segmentation was performed using the Nuclear envelope segmentation system (NesSys) (Blin et al 2019) using LaminB1 signal. Ten of the 19 segmented embryos were manually edited in PickCells to correct for segmentation errors.

#### High throughput single cell quantification

Image analysis on images from a high content confocal microscopy platform, including 2D segmentation and signal quantification, was carried out in the Signals Image Artist environment (Revvity Signals Software). <u>Neighbour identification</u> was carried out in PickCells by Delaunay triangulation using nuclei centroids as input. For embryos, the epiblast was manually labelled in PickCells and only the epiblast nuclei were used as input for neighbour identification (see Fig 2A).

Nuclei to neighbour edges and individual nuclei features, were exported to R for all subsequent analysis including neighbour kernel smoothing, epiblast normalisation procedure, and pseudospace identification.

#### Signal correction and normalisation

All subsequent analysis was performed with custom scripts in R. We first used linear unmixing to correct for signal bleed through between the TBXT and TBX6 channels. We also corrected for the loss of fluorescence intensity across the depth of the image due to light scattering effects. Next, raw values of mean signal intensity per nuclei were normalised by the formula (signal - background)/ (max signal - background). The max signal was chosen as the 99th percentile to exclude hot pixels and signal coming from staining artefacts such as cell debris and antibody aggregates. The background signal was overrepresented in the image and was estimated by finding the highest peak in the signal histogram. Nuclei were classified as positive or negative by gating the populations manually after combining all the replicates per experiment.

#### Nuclear intensities smoothing

TF signal for each nuclei was iteratively smoothed by taking the average TF signal of the nuclei and its neighbours. This new average (AvTF+1) was then used in another round, where the average AvTF signal of a nucleus was calculated (AvTF+2). This was repeated 10 times (AvTF+10) for each TF measured to spatially smooth the TF signal. After which, the log (AvTF+10) signal was used as an input for PCA dimension reduction.

#### Pseudotime construction and NMP gating

The pseudo-space route from Neural to NMP to Mesoderm was identified using the Slingshot package (Street et al., 2018) in PCA1 and PCA2 space. NMP region gates in pseudospace were ascertained by scanning gate values to best fit the bi-fated region in four somite pair embryos, which is then applied to all wild type embryo stages. Gates in pseudospace used to isolate NMP ROIs in ex vivo cultured embryos were identified by i) the average pseudospace of node streak border nuclei and ii) the average pseudospace values when mean population TBX6 rises above background along the pseudospace axis.

The coefficient of variation of TF signal intensity for a nuclei and its neighbours defined as the cell niche (*X* = {*x_1_, x_2_, x_3_* … *x*_n_}), defined as the standard deviation (σ_*X*_) normalised by the mean (*X*) as below:

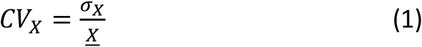

The neighbour ratio (NR) is calculated as the ratio of a nuclei’s (C) mean TF signal value compared to the average of its neighbouring nuclei (*N* = {n, n*_2_*, n*_3_* … n_n_}) defined as below:

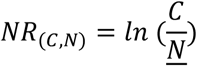

TF values were simulated using the rnorm function in the compositions package in R. This function generates random numbers within a normal distribution, defined by a standard deviation and mean provided. In this case, the inputs for rnorm were calculated as in CV, using the standard deviation and mean of a nucleus’s TF expression and the TF expression of its neighbours.

#### Data representation

Figures were created with custom scripts in R and MATLAB. Tools used to plot data include violin plots [63] (Kenny and Shoen et al 2021). Colour maps include the scientific colour map package scico (Crameri 2018, Scientific colour maps. Zenodo. doi: 10.5281/ZENODO.1243909). Additional details of image analysis methods are provided in Supplemental Methods.

## Supplemental Information

### 8 supplemental figures

**Fig S1.**
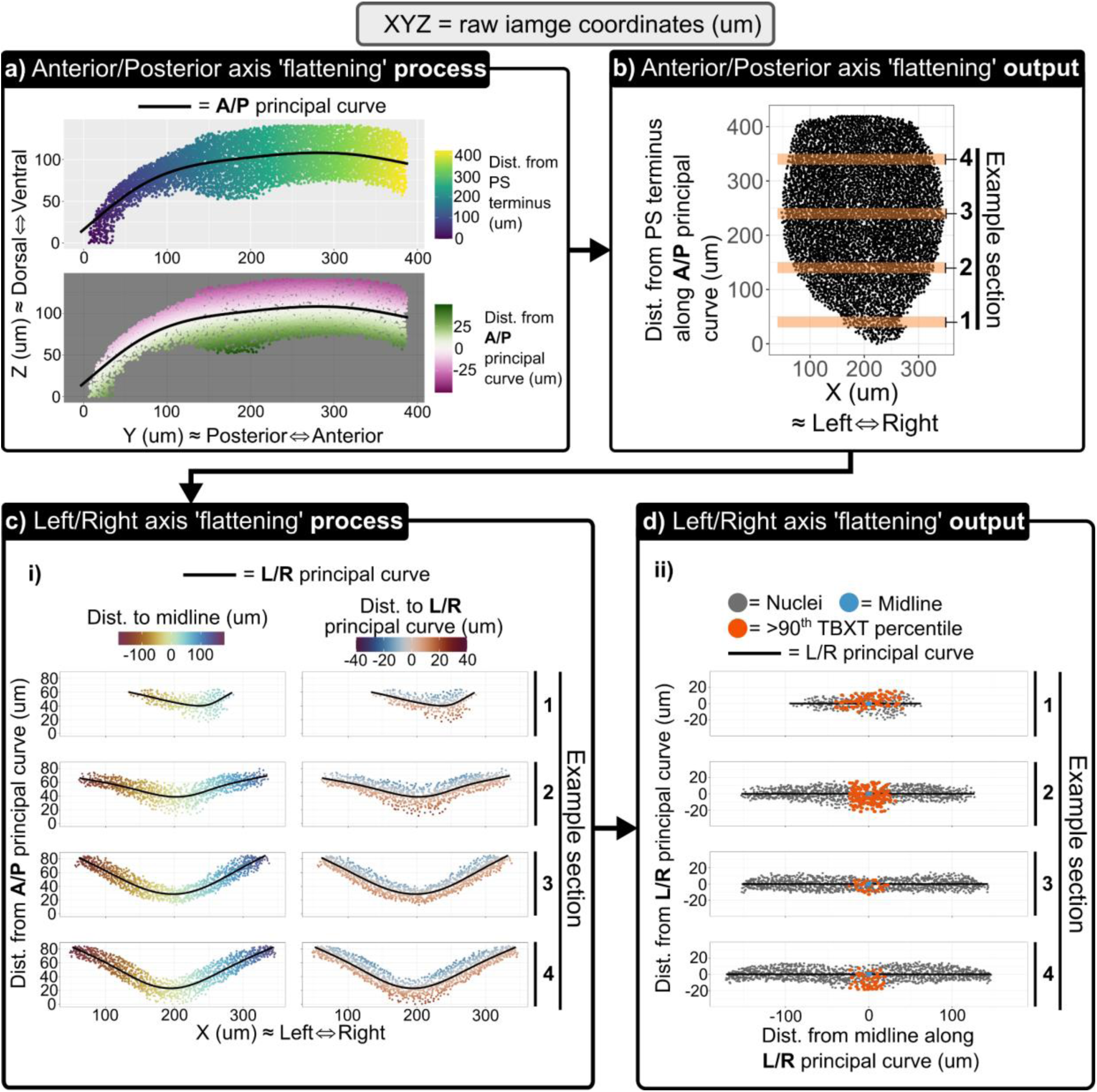
3D Manifold projection and alignment Example of the algorithm modules on a segmented six somite pair embryo with the epiblast manually labelled. Points represent nuclei centroids and XYZ refer to coordinates in the raw image planes. In this process (**a)** first the position of each nuclei in the anterior posterior axis calculated relative to the nearest point on a principal curve in the Y and Z planes, which roughly correspond to the anterior/posterior (A/P) and dorsal/ventral axes respectively. The dorsal/ventral axis is initially estimated as the distance to the nearest point on this A/P principal curve. The output of this **(b)** is carried to the next step (**b)** to identify the position along the left/right (L/R) and the dorsal/ventral (D/V) axis. In this step, sections of the epiblast parallel to the new A/P axis are isolated, four example sections are shown. In **(c)** Relative positions of nuclei to a principal curve in the X plane (roughly corresponds to the left/right axis) and the estimated D/V axis [distance to the A/P principal curve from **(a)** are calculated to identify the L/R and D/V axes. **(d)** The distances of each nuclei along the principal curve are normalised to the ‘midline’, which is the average L/R position of the 90^th^ percentile of spatially smoothed TBXT values.

**Fig S2.**
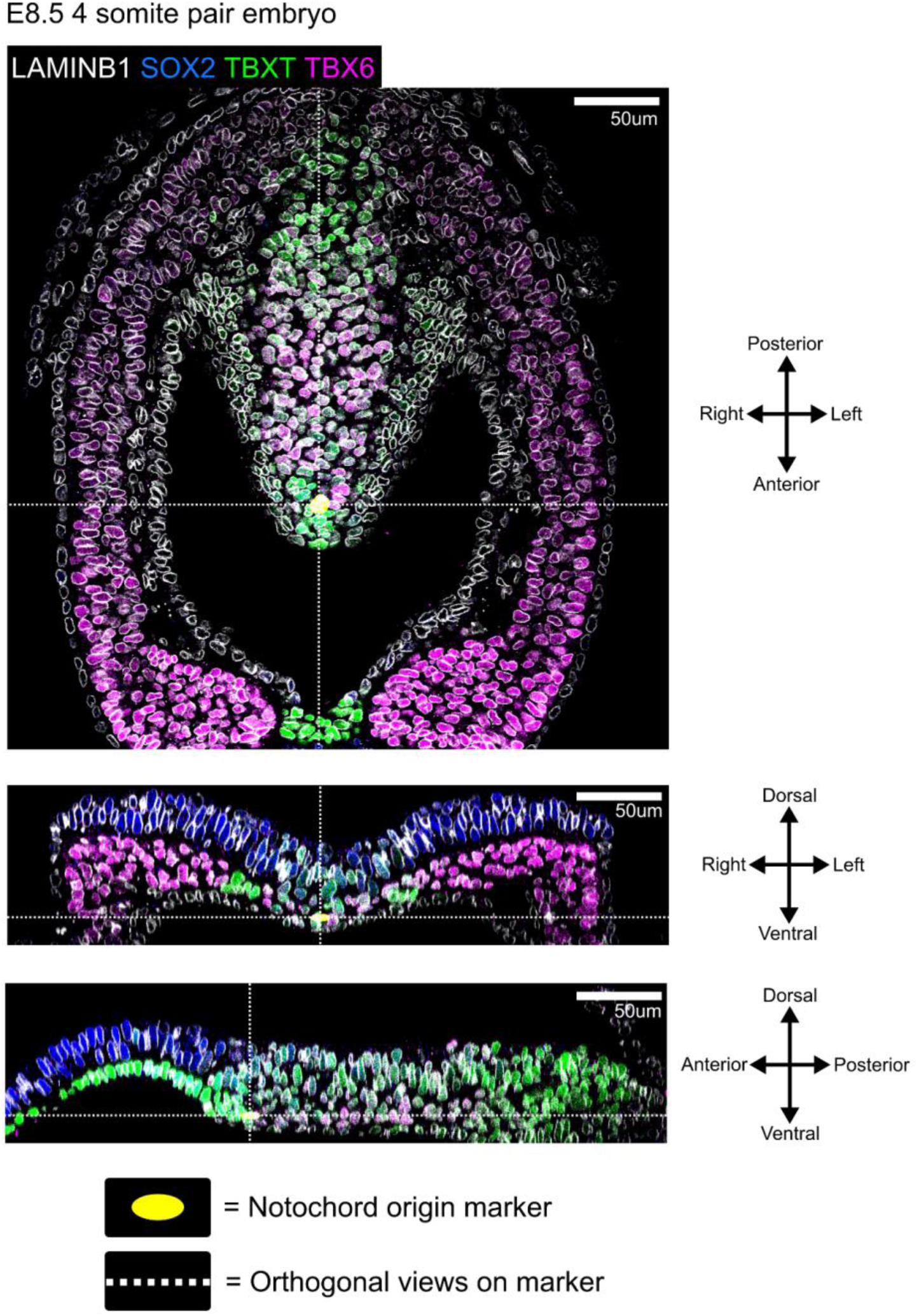
Locating embryo landmarks Example of the manual marking of the posteriormost end of the notochord in a confocal stack of four somite pair embryo stained for LaminB1, SOX2, TBXT, and TBX6, with orthogonal views.

**Fig S3.**
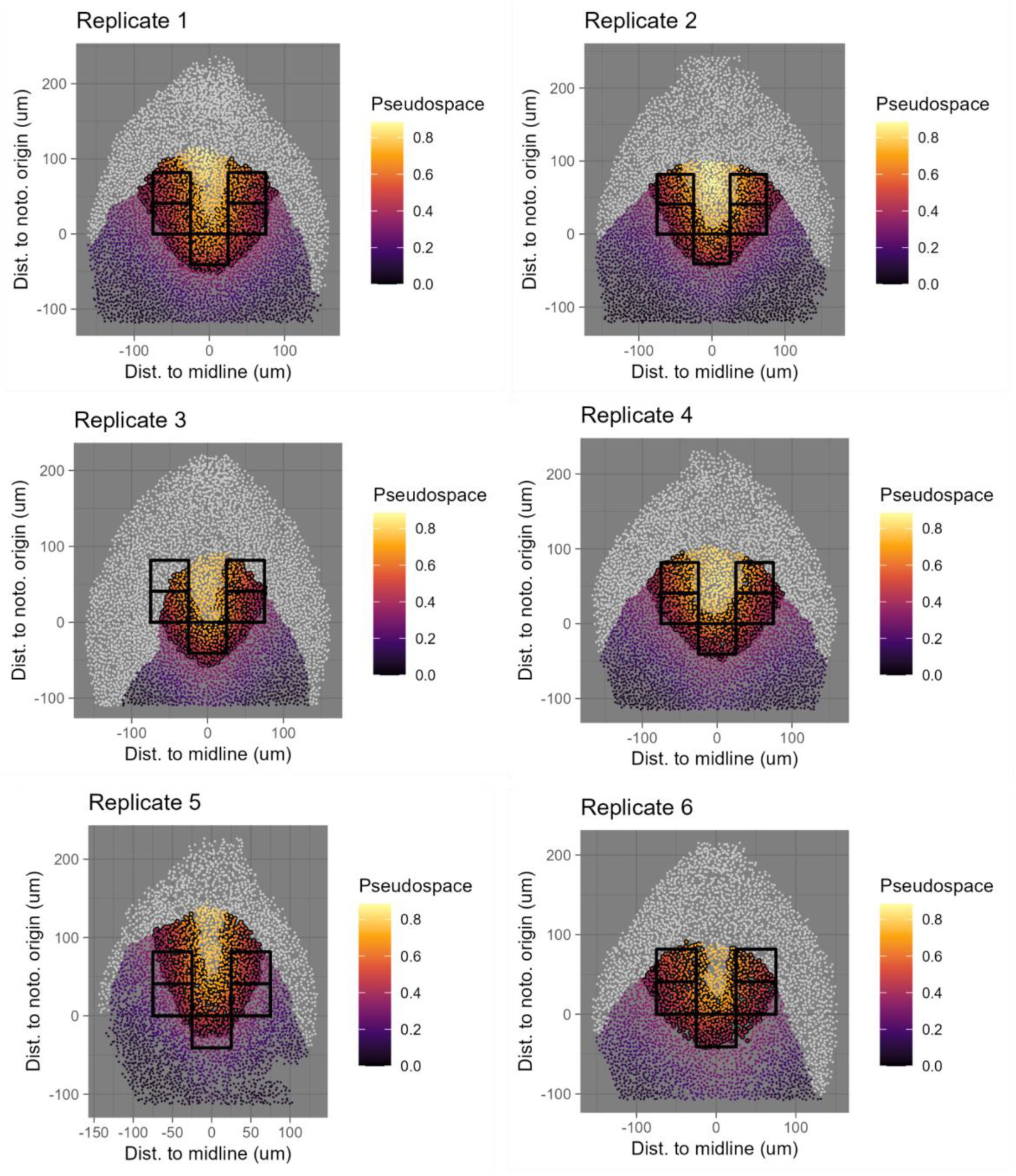
Mapping putative NMP regions Four somite pair embryo epiblast projections showing manual best-fits of pseudospace gates to bifated regions (in boxes). Showing left/right asymmetry in replicate three and non-specific bi-fated region labelling in replicate 5.

**Fig S4.**
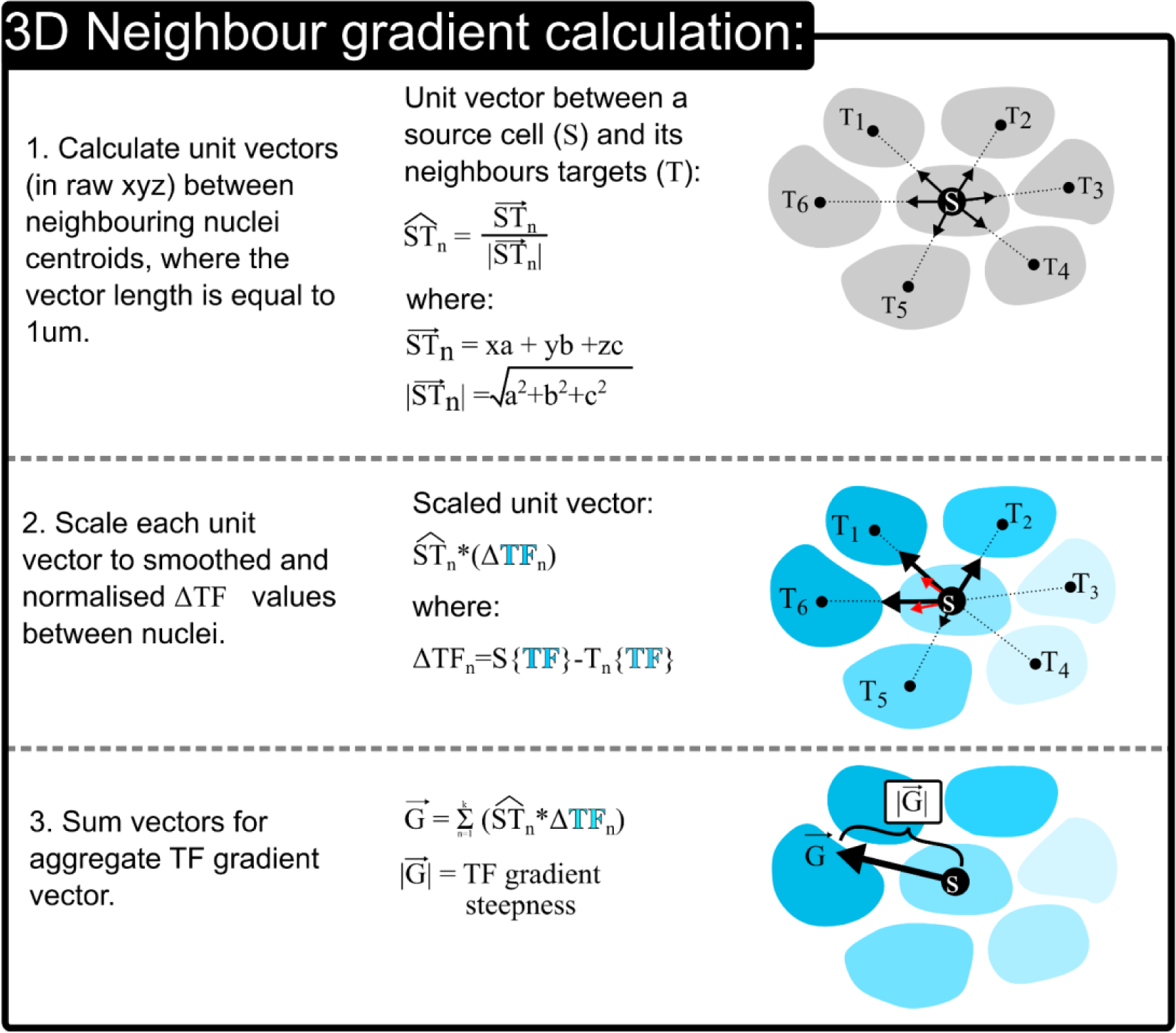
3D gradient neighbour calculation method An explanation of the method for calculating local gradient direction and steepness

**Fig S5.**
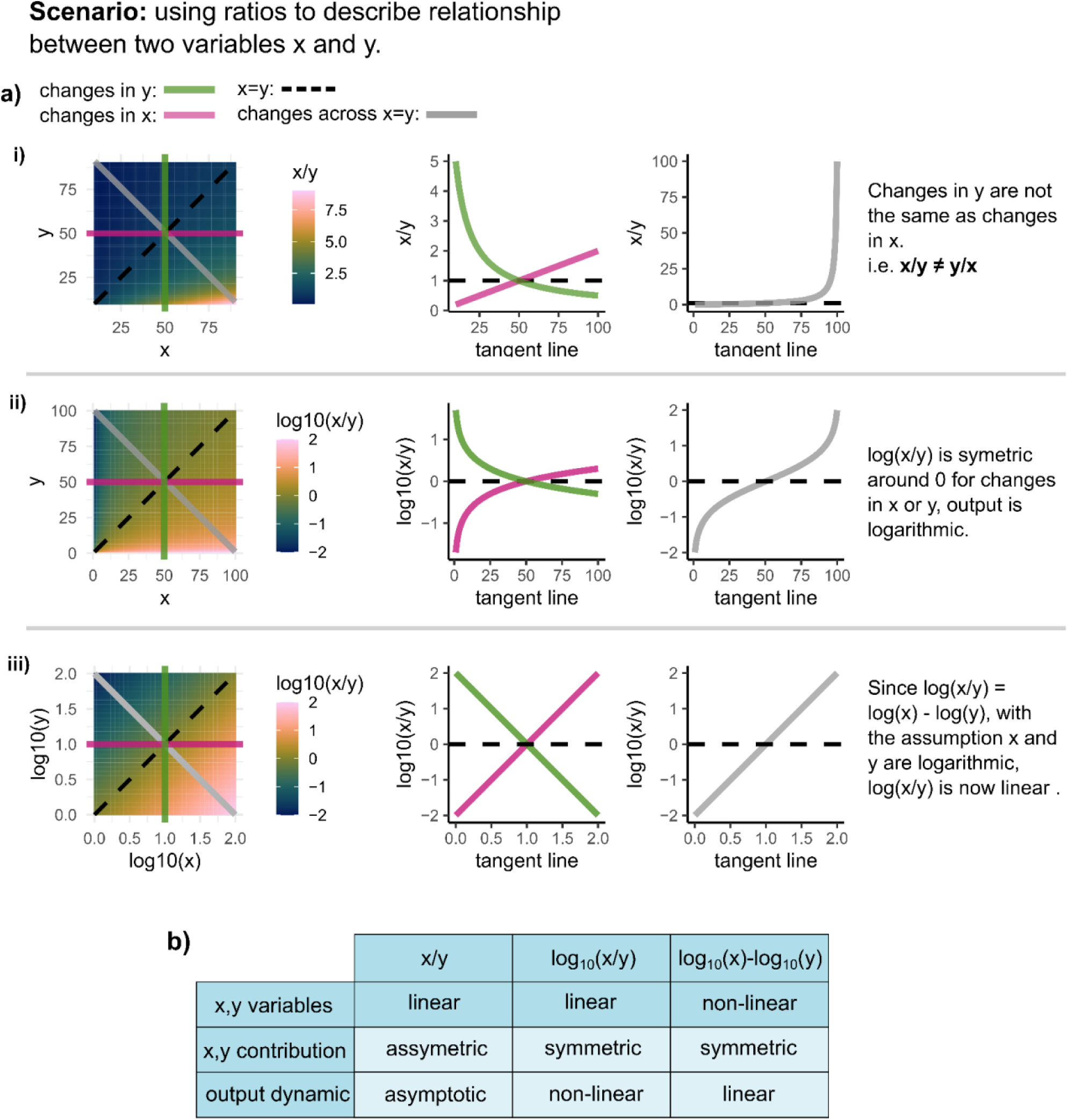
Using ratios to describe relationship between two variables **a)** Outline of scenarios to explain how log(x/y) is suitable to describe the relationship between two variables. **i)** In this scenario, x/y does not equal y/x, and y has logarithmic and asymmetric influence on the output of the calculation. **ii)** using log(x/y) makes the calculation symmetric around 0, but the output is now non-linear. **iii)** If the variables x and y can be assumed to be non-linear and a log transformation of x and y is appropriate, then the relationships between log(x) and log(y) with log(x/y) is now linear due to the log law log(x/y) = log(x) – log(y). **b)** Summary of the scenarios and dynamic of the calculation output.

**Fig S6.**
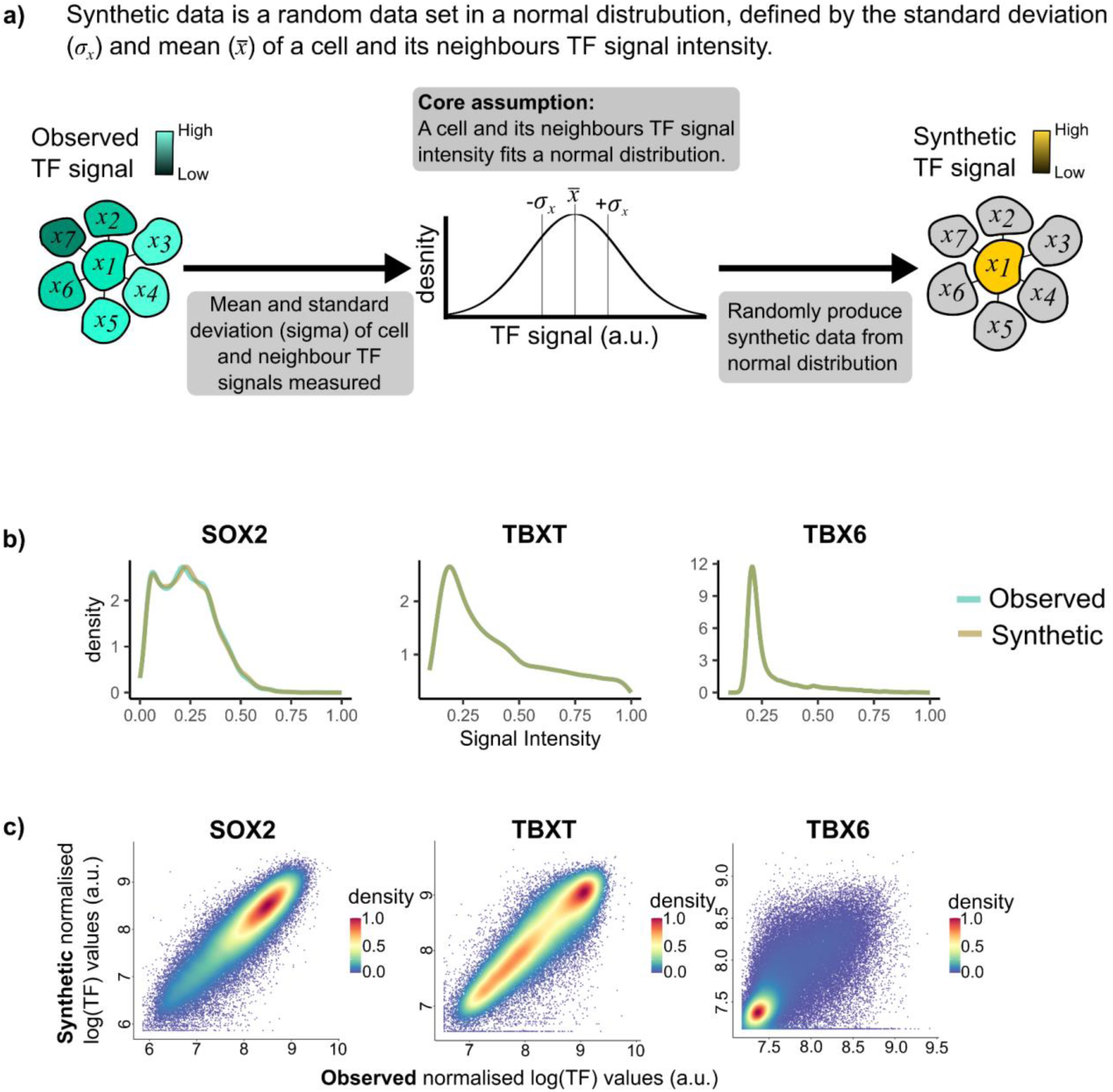
Synthetic data **a)** Process to produce synthetic data. **b)** Comparison of global population distribution for observed and synthetic SOX2, TBXT, and TBX6 signal intensity values. One four somite pair embryo dataset shown. **c)** Density plots to compare observed and synthetic TF values of individual nuclei, showing close associations of SOX2 and TBXT, but less accurate TBX6 associations. All embryos shown n=19.

**Fig S7.**
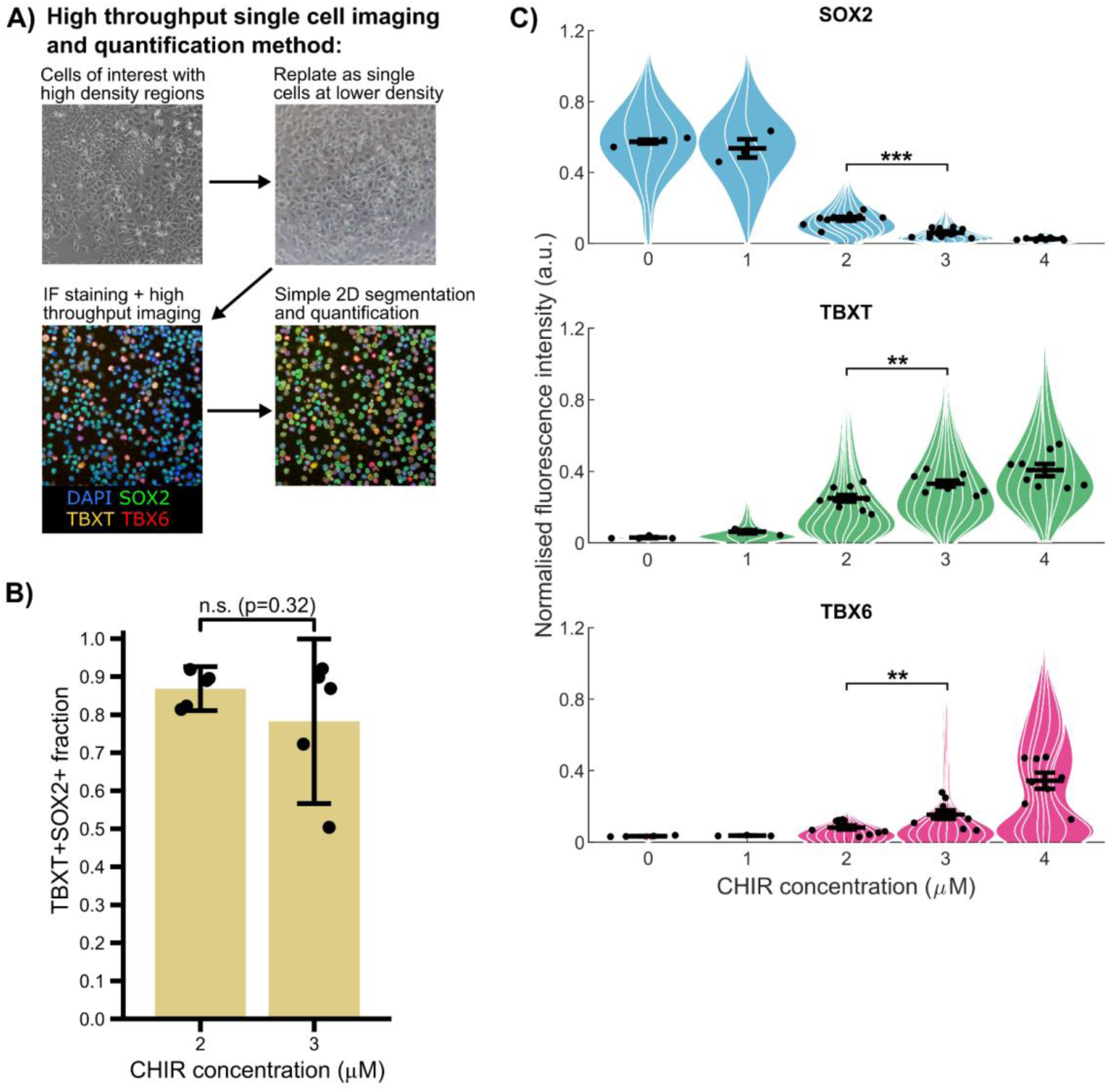
Human ESC to NMP differentiation characterisation and efficiency. **A)** High throughput single cell quantification method used to complement high resolution imaging (used in spatial neighbour analysis) by boosting sample sizes and N numbers. hNMP monolayers typically form dense structures which are difficult to segment and require time-consuming high-resolution images. To address this, cells are replated as single cells at a lower density and then quickly fixed to perform IF and stain for hNMP markers and DAPI. These are imaged on high imaging content platforms and present a simple challenge for single cell segmentation methods even at low resolutions. **B)** Differentiation of human MShef7 to hNMPs with 20ng bFGF and 2uM or 3uM CHIR (as described in figure 6a) both produce high proportions of TBXT+SOX2+ populations with no statistically significant difference between the groups. (n=5). **C)** CHIR titration of the hNMP differentiation protocol shows the influence of CHIR on SOX2, TBXT, and TBX6, where 3μM CHIR produces a population with higher TBXT/TBX6 and lower SOX2 than 2μM (n=9). Points on violin superplot and barplot show mean per replicate. All error bars indicate confidence intervals of 0.95. Statistical tests performed by one way ANOVA, ** = p<0.01, *** = p<0.001.

**Fig S8.**
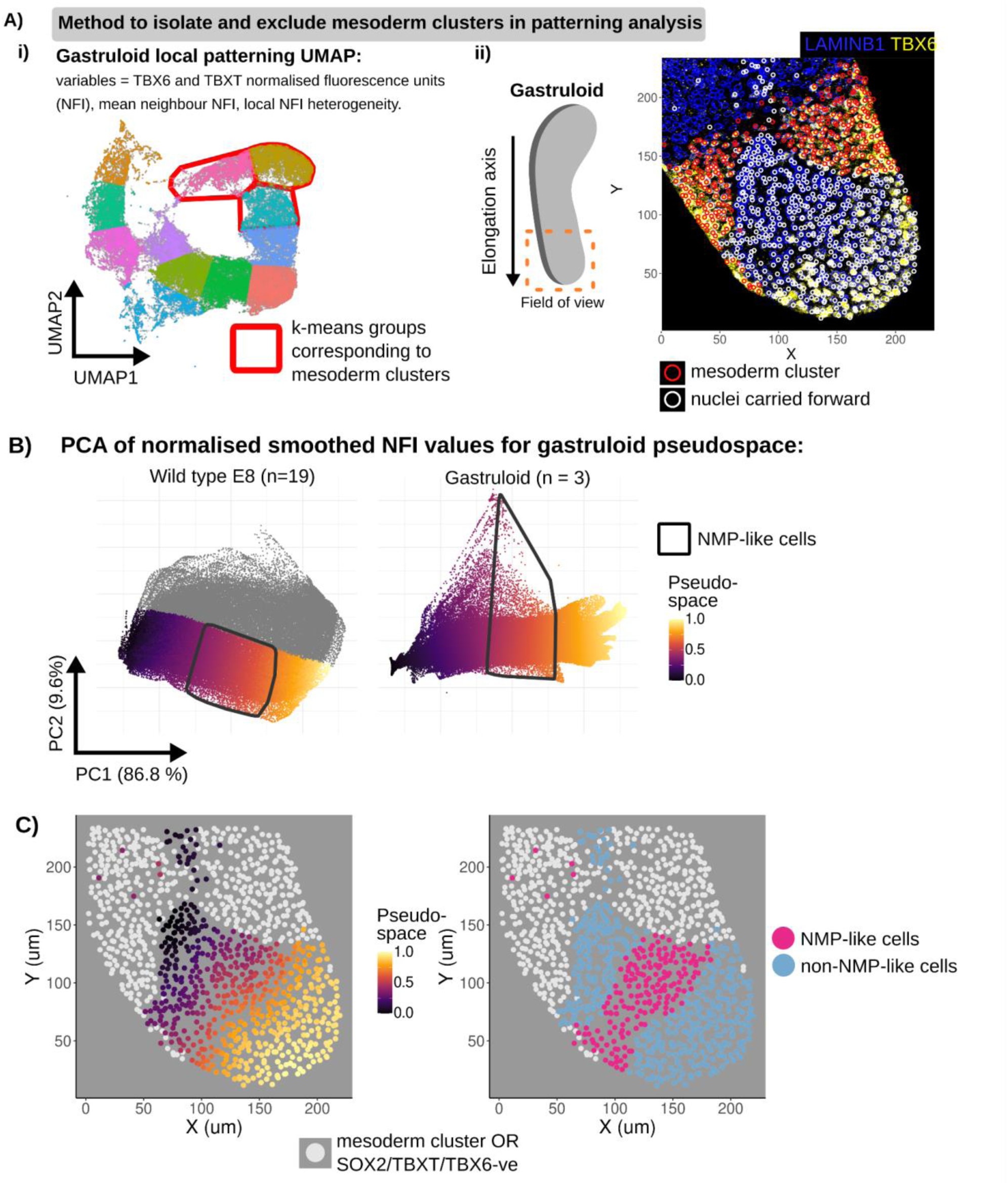
Methodology to identify NMP-like profiles in gastruloids using pseudospace A. Process in identifying and excluding paraxial mesoderm-like clusters in gastruloids. **i)** UMAP of TBXT/TBX6 patterning variables; namely normalised fluorescence units (NFI), mean neighbour NFI, local NFI heterogeneity (coefficient of variation). K-means clustering identifies clusters which when **(ii)** mapped onto the gastruloid localises with paraxial mesoderm-like clusters. Graphic showing field of view for IF image and overlay of mesoderm-clusters and TF +ve cells. Mesoderm clusters are excluded in downstream analysis. B. Similar in figure 3, slingshot pseudotime tools are used in a PCA dimension reduction of smoothed normalised fluorescence units in gastruloids to identify pseudospace. NMP-like cells (NMPLC) are identified in gastruloids using pseudospace gates characterised E8 embryos. C. Pseudospace values and NMPLC mapped onto the 10um section of a gastruloid using nuclei centroids. NMPLCs can be found not at the tip, but just ‘anterior’ of the tip in the elongation axis.

#### 1 supplemental methods document giving details of algorithms

## Supplemental methods: details of algorithms

### Epiblast manifold projection and alignment algorithm steps

1. The raw XYZ coordinates of an individual embryo epiblast are manually oriented such that the Anterior Posterior axis aligns with the Y axis, such that the epiblast is as close to symmetric and the Z axis is as flat as possible, creating new XYZ coordinates.
2. Each cell’s position along an interpolated principle curve (as measured 0->*n* in um) in the new YZ plane is using found the slingshot package (Street et al., 2018). This metric is used herein as the anterior-posterior (AP) axis position.
3. Starting from 0 in a moving window, cells +/- 20-30um* along from this AP position are isolated. This is the start of the loop; the AP position will increase in steps of ∼5um* and run through to the terminal nuclei of the epiblast.
4. In the XZ plane, each cell’s position along the epiblast curve is found using the slingshot package (as measured 0->*n* in um). This is termed the left/right (LR) position value. The direction of position 0->n along the curve is ensured to orient from left to right.
5. The cells expressing the top 10%* levels of T are identified, and the average LR value of these cells is determined as the midline, not any individual cell.
6. The LR midline value is subtracted from all other cell’s LR value, in doing so normalising the position of each cell to the midline. Cells to the left of the midline have negative LR values, and to the right positive LR values.
7. After this normalisation, the distance from the midline as a percentage from the epiblast edge is calculated. First, the LR values for cells to the left of the midline are divided by the LR value furthest cell on the right side (identified by containing a negative LR value), and similarly for the left side (identified by containing a positive LR value).
8. This information is stored and loops back to step 3, where the next AP position is +∼5um (a parameter for optimisation) from the previous subset position. The small increase in AP position results in each cell’s LR position being calculated multiple times and a moving window.
9. After the entire AP axis has been run through, the multiple values of LR position and relative LR position for each cell has been calculated, among other metrics. The average of these values from every loop is carried forward. * = parameters optimised per dataset.

### Epiblast shape registration method algorithm steps

1. First, the posterior nascent notochord is manually labelled in each epiblast in PickCells, and the equivalent AP position of this is subtracted from each cell’s AP value. Such that position 0 is the start of the notochord.
2. To normalise the length of the primitive streak, each cell’s new notochord normalised AP value is divided by the posterior most value, creating a relative position along the primitive streak from the Notochord. This relative AP value is carried forward.
3. Next, the LR position is either maintained as right negative and left positive values are converted to positive values, effectively digitally folding at the midline to combine LR sides of the epiblast.
4. Starting from the posterior end in a moving window, cells from all epiblasts +/- 0.05* relative AP values from the starting position (S) are isolated.
5. The relative LR value is multiplied
6. Then, the process loops back to step 4, where the AP position is increased by a step of 0.015 as a moving window. Again, calculating multiple values of renormalised LR values for each cell. The loop finishes at the posterior most cell’s AP position.
7. After the process has run through, the average of the renormalised LR values for each cell are used as the final normalised LR position. * = parameters optimised per experiment.

### Neighbour Smoothing and Neural-NMP-Mesoderm pseudo-space

TF signal for each nuclei was iteratively smoothed by taking the average TF signal of the nuclei and its neighbours. This new average (AvTF+1) was then used in another round, where the average AvTF signal of a nucleus was calculated (AvTF+2). This was repeated 10 times (AvTF+10) for each TF measured to spatially smooth the TF signal. After which, the log(AvTF+10) signal was used as an input for PCA dimension reduction. The pseudo-space route from Neural to NMP to Mesoderm was identified using the Slingshot package (Street et al., 2018) in PCA1 and PCA2 space. NMP region gates in pseudospace were ascertained by scanning gate values to best fit the bi-fated region in four somite pair embryos, which is then applied to all wild type embryo stages. Gates in pseudospace used to isolate NMP ROIs in ex *vivo* cultured embryos were identified by i) the average pseudospace of node streak border nuclei and ii) the average pseudospace values when mean population TBX6 rises above background along the pseudospace axis. STREET, K., RISSO, D., FLETCHER, R. B., DAS, D., NGAI, J., YOSEF, N., PURDOM, E. & DUDOIT, S. 2018. Slingshot: cell lineage and pseudotime inference for single-cell transcriptomics. *BMC Genomics,* 19, 477.

## Notes

### Competing Interest Statement

The authors have declared no competing interest.

## References

1. Briscoe J, Small S. Morphogen rules: design principles of gradient-mediated embryo patterning. Development. 2015;142: 3996–4009.

2. Berg S, Kutra D, Kroeger T, Straehle CN, Kausler BX, Haubold C, et al. Ilastik: Interactive machine learning for (bio)image analysis. Nat Methods. 2019;16: 1226–1232.

3. Blin G, Sadurska D, Portero Migueles R, Chen N, Watson JA, Lowell S. Nessys: A new set of tools for the automated detection of nuclei within intact tissues and dense 3D cultures. PLoS Biol. 2019;17: e3000388.

4. Weigert M, Schmidt U, Haase R, Sugawara K, Myers G. Star-convex polyhedra for 3D object detection and segmentation in microscopy. Proceedings of the IEEE/CVF winter conference on applications of computer vision. 2020. pp. 3666–3673.

5. Eschweiler D, Smith RS, Stegmaier J. Robust 3d Cell Segmentation: Extending The View Of Cellpose. 2022 IEEE International Conference on Image Processing (ICIP). 2022. pp. 191–195.

6. Moen E, Bannon D, Kudo T, Graf W, Covert M, Van Valen D. Deep learning for cellular image analysis. Nat Methods. 2019;16: 1233–1246.

7. Pachitariu M, Stringer C. Cellpose 2.0: how to train your own model. Nat Methods. 2022;19: 1634– 1641.

8. Forsyth JE, Al-Anbaki AH, de la Fuente R, Modare N, Perez-Cortes D, Rivera I, et al. IVEN: A quantitative tool to describe 3D cell position and neighbourhood reveals architectural changes in FGF4-treated preimplantation embryos. PLoS Biol. 2021;19: e3001345.

9. Fischer SC, Schardt S, Lilao-Garzón J, Muñoz-Descalzo S. The salt-and-pepper pattern in mouse blastocysts is compatible with signaling beyond the nearest neighbors. iScience. 2023;26: 108106.

10. Henrique D, Schweisguth F. Mechanisms of Notch signaling: a simple logic deployed in time and space. Development. 2019;146. doi:10.1242/dev.172148

11. Johnston RJ Jr, Desplan C. Stochastic mechanisms of cell fate specification that yield random or robust outcomes. Annu Rev Cell Dev Biol. 2010;26: 689–719.

12. Petrovic J, Formosa-Jordan P, Luna-Escalante JC, Abelló G, Ibañes M, Neves J, et al. Ligand-dependent Notch signaling strength orchestrates lateral induction and lateral inhibition in the developing inner ear. Development. 2014;141: 2313–2324.

13. Linker C, De Almeida I, Papanayotou C, Stower M, Sabado V, Ghorani E, et al. Cell communication with the neural plate is required for induction of neural markers by BMP inhibition: evidence for homeogenetic induction and implications for Xenopus animal cap and chick explant assays. Dev Biol. 2009;327: 478–486.

14. Chen C-C, Wang L, Plikus MV, Jiang TX, Murray PJ, Ramos R, et al. Organ-level quorum sensing directs regeneration in hair stem cell populations. Cell. 2015;161: 277–290.

15. Gurdon JB, Lemaire P, Kato K. Community effects and related phenomena in development. Cell. 1993;75: 831–834.

16. Nemashkalo A, Ruzo A, Heemskerk I, Warmflash A. Morphogen and community effects determine cell fates in response to BMP4 signaling in human embryonic stem cells. Development. 2017;144: 3042– 3053.

17. Lowell S. You should always keep in touch with your friends: Community effects in biology. Nat Rev Mol Cell Biol. 2020;21: 568–569.

18. Luengo Hendriks CL, Keränen SVE, Fowlkes CC, Simirenko L, Weber GH, DePace AH, et al. Three-dimensional morphology and gene expression in the Drosophila blastoderm at cellular resolution I: data acquisition pipeline. Genome Biol. 2006;7: R123.

19. Guirao B, Rigaud SU, Bosveld F, Bailles A, López-Gay J, Ishihara S, et al. Unified quantitative characterization of epithelial tissue development. Elife. 2015;4. doi:10.7554/eLife.08519

20. Heller D, Hoppe A, Restrepo S, Gatti L, Tournier AL, Tapon N, et al. EpiTools: An Open-Source Image Analysis Toolkit for Quantifying Epithelial Growth Dynamics. Dev Cell. 2016;36: 103–116.

21. Stadler T, Skylaki S, D Kokkaliaris K, Schroeder T. On the statistical analysis of single cell lineage trees. J Theor Biol. 2018;439: 160–165.

22. Toth T, Balassa T, Bara N, Kovacs F, Kriston A, Molnar C, et al. Environmental properties of cells improve machine learning-based phenotype recognition accuracy. Sci Rep. 2018;8: 10085.

23. Hicks DG, Speed TP, Yassin M, Russell SM. Maps of variability in cell lineage trees. PLoS Comput Biol. 2019;15: e1006745.

24. Shah G, Thierbach K, Schmid B, Waschke J, Reade A, Hlawitschka M, et al. Multi-scale imaging and analysis identify pan-embryo cell dynamics of germlayer formation in zebrafish. Nat Commun. 2019;10: 5753.

25. Gómez HF, Dumond MS, Hodel L, Vetter R, Iber D. 3D cell neighbour dynamics in growing pseudostratified epithelia. Elife. 2021;10. doi:10.7554/eLife.68135

26. Calvo L, Ronshaugen M, Pettini T. smiFISH and embryo segmentation for single-cell multi-gene RNA quantification in arthropods. Commun Biol. 2021;4: 352.

27. Strauss S, Runions A, Lane B, Eschweiler D, Bajpai N, Trozzi N, et al. Using positional information to provide context for biological image analysis with MorphoGraphX 2.0. Elife. 2022;11. doi:10.7554/eLife.72601

28. Summers HD, Wills JW, Rees P. Spatial statistics is a comprehensive tool for quantifying cell neighbor relationships and biological processes via tissue image analysis. Cell Rep Methods. 2022;2: 100348.

29. Blanchard GB, Fletcher AG, Schumacher LJ. The devil is in the mesoscale: Mechanical and behavioural heterogeneity in collective cell movement. Semin Cell Dev Biol. 2019;93: 46–54.

30. Wong MD, van Eede MC, Spring S, Jevtic S, Boughner JC, Lerch JP, et al. 4D atlas of the mouse embryo for precise morphological staging. Development. 2015;142: 3583–3591.

31. McDole K, Guignard L, Amat F, Berger A, Malandain G, Royer LA, et al. In Toto Imaging and Reconstruction of Post-Implantation Mouse Development at the Single-Cell Level. Cell. 2018;175: 859– 876.e33.

32. Wymeersch FJ, Wilson V, Tsakiridis A. Understanding axial progenitor biology in vivo and in vitro. Development. 2021;148. doi:10.1242/dev.180612

33. Martins-Costa C, Wilson V, Binagui-Casas A. Neuromesodermal specification during head-to-tail body axis formation. Curr Top Dev Biol. 2024;159: 232–271.

34. Romanos M, Allio G, Roussigné M, Combres L, Escalas N, Soula C, et al. Cell-to-cell heterogeneity in Sox2 and Bra expression guides progenitor motility and destiny. Elife. 2021;10. doi:10.7554/eLife.66588

35. Hatakeyama Y, Saito N, Mii Y, Takada R, Shinozuka T, Takemoto T, et al. Intercellular exchange of Wnt ligands reduces cell population heterogeneity during embryogenesis. Nat Commun. 2023;14: 1924.

36. Chapman D, Papaioannou VE. Three neural tubes in mouse embryos with mutations in the T-box gene Tbx6. Nature. 1998;391: 695–697.

37. Chapman D, Cooper-Morgan A, Harrelson Z, Papaioannou VE. Critical role for Tbx6 in mesoderm specification in the mouse embryo. Mech Dev. 2003;120: 837–847.

38. Takemoto T, Uchikawa M, Yoshida M, Bell DM, Lovell-Badge R, Papaioannou VE, et al. Tbx6-dependent Sox2 regulation determines neural or mesodermal fate in axial stem cells. Nature. 2011;470: 394–398.

39. Cooper F, Souilhol C, Haston S, Gray S, Boswell K, Gogolou A, et al. Notch signalling influences cell fate decisions and HOX gene induction in axial progenitors. Development. 2024;151. doi:10.1242/dev.202098

40. Cambray N, Wilson V. Two distinct sources for a population of maturing axial progenitors. Development. 2007;134: 2829–2840.

41. Wymeersch FJ, Huang Y, Blin G, Cambray N, Wilkie R, Wong FCK, et al. Position-dependent plasticity of distinct progenitor types in the primitive streak. Elife. 2016;5: e10042.

42. Binagui-Casas A, Dias A, Guillot C, Metzis V, Saunders D. Building consensus in neuromesodermal research: Current advances and future biomedical perspectives. Curr Opin Cell Biol. 2021;73: 133–140.

43. Street K, Risso D, Fletcher RB, Das D, Ngai J, Yosef N, et al. Slingshot: cell lineage and pseudotime inference for single-cell transcriptomics. BMC Genomics. 2018;19: 477.

44. Gouti M, Delile J, Stamataki D, Wymeersch FJ, Huang Y, Kleinjung J, et al. A Gene Regulatory Network Balances Neural and Mesoderm Specification during Vertebrate Trunk Development. Developmental Cell. 2017. pp. 243–261.e7. doi:10.1016/j.devcel.2017.04.002

45. Wind M, Tsakiridis A. In Vitro Generation of Posterior Motor Neurons from Human Pluripotent Stem Cells. Curr Protoc. 2021;1: e244.

46. van den Brink SC, Baillie-Johnson P, Balayo T, Hadjantonakis A-K, Nowotschin S, Turner DA, et al. Symmetry breaking, germ layer specification and axial organisation in aggregates of mouse embryonic stem cells. Development. 2014;141: 4231–4242.

47. van den Brink SC, Alemany A, van Batenburg V, Moris N, Blotenburg M, Vivié J, et al. Single-cell and spatial transcriptomics reveal somitogenesis in gastruloids. Nature. 2020;582: 405–409.

48. Warchol S, Krueger R, Nirmal AJ, Gaglia G, Jessup J, Ritch CC, et al. Visinity: Visual Spatial Neighborhood Analysis for Multiplexed Tissue Imaging Data. IEEE Trans Vis Comput Graph. 2023;29: 106–116.

49. Blin G. Quantitative developmental biology in vitro using micropatterning. Development. 2021;148. doi:10.1242/dev.186387

50. Haertter D, Wang X, Fogerson SM, Ramkumar N, Crawford JM, Poss KD, et al. DeepProjection: specific and robust projection of curved 2D tissue sheets from 3D microscopy using deep learning. Development. 2022;149. doi:10.1242/dev.200621

51. Wymeersch FJ, Skylaki S, Huang Y, Watson JA, Economou C, Marek-Johnston C, et al. Transcriptionally dynamic progenitor populations organised around a stable niche drive axial patterning. Development. 2019;146. doi:10.1242/dev.168161

52. Corson LB, Yamanaka Y, Lai K-MV, Rossant J. Spatial and temporal patterns of ERK signaling during mouse embryogenesis. Development. 2003;130: 4527–4537.

53. Olivera-Martinez I, Schurch N, Li RA, Song J, Halley PA, Das RM, et al. Major transcriptome re-organisation and abrupt changes in signalling, cell cycle and chromatin regulation at neural differentiation in vivo. Development. 2014;141: 3266–3276.

54. Morgani SM, Saiz N, Garg V, Raina D, Simon CS, Kang M, et al. A Sprouty4 reporter to monitor FGF/ERK signaling activity in ESCs and mice. Dev Biol. 2018;441: 104–126.

55. Morgani SM, Hadjantonakis A-K. Signaling regulation during gastrulation: Insights from mouse embryos and in vitro systems. Curr Top Dev Biol. 2020;137: 391–431.

56. Akai J, Halley PA, Storey KG. FGF-dependent Notch signaling maintains the spinal cord stem zone. Genes Dev. 2005;19: 2877–2887.

57. Perez-Carrasco R, Guerrero P, Briscoe J, Page KM. Intrinsic Noise Profoundly Alters the Dynamics and Steady State of Morphogen-Controlled Bistable Genetic Switches. PLoS Comput Biol. 2016;12: e1005154.

58. Fulton T, Spiess K, Thomson L, Wang Y, Clark B, Hwang S, et al. Cell Rearrangement Generates Pattern Emergence as a Function of Temporal Morphogen Exposure. 2021. doi:10.1101/2021.02.05.429898

59. Malaguti M, Portero Migueles R, Annoh J, Sadurska D, Blin G, Lowell S. SyNPL: Synthetic Notch pluripotent cell lines to monitor and manipulate cell interactions in vitro and in vivo. Development. 2022;149. doi:10.1242/dev.200226

60. Lebek T, Malaguti M, Elfick A, Lowell S. PUFFFIN: A novel, ultra-bright, customisable, single-plasmid system for labelling cell neighbourhoods. BioRxiv. 2023. doi: 10.1101/2023.09.06.556381

61. Malaguti M, Lebek T, Blin G, Lowell S. Enabling neighbour labelling: using synthetic biology to explore how cells influence their neighbours. Development. 2024;151. doi:10.1242/dev.201955

62. Smith AG, Hooper ML. Buffalo rat liver cells produce a diffusible activity which inhibits the differentiation of murine embryonal carcinoma and embryonic stem cells. Dev Biol. 1987;121: 1–9.

63. Kenny M, Schoen I. Violin SuperPlots: visualizing replicate heterogeneity in large data sets. Mol Biol Cell. 2021;32: 1333–1334.

